# Laminar Neural Dynamics of Auditory Evoked Responses: Computational Modeling of Local Field Potentials in Auditory Cortex of Non-Human Primates

**DOI:** 10.1101/2022.12.21.521407

**Authors:** Vincent S.C. Chien, Peng Wang, Burkhard Maess, Yonatan Fishman, Thomas R. Knösche

## Abstract

Evoked neural responses to sensory stimuli have been extensively investigated in humans and animal models both to enhance our understanding of brain function and to aid in clinical diagnosis of neurological and neuropsychiatric conditions. Recording and imaging techniques such as electroencephalography (EEG), magnetoencephalography (MEG), local field potentials (LFPs), and calcium imaging provide complementary information about different aspects of brain activity at different spatial and temporal scales. Modeling and simulations provide a way to integrate these different types of information to clarify underlying neural mechanisms.

In this study, we aimed to shed light on the neural dynamics underlying auditory evoked responses by fitting a rate-based model to LFPs recorded via multi-contact electrodes which simultaneously sampled neural activity across cortical laminae. Recordings included neural population responses to best-frequency (BF) and non-BF tones at four representative sites in primary auditory cortex (A1) of awake monkeys. The model considered major neural populations of excitatory, parvalbumin-expressing (PV), and somatostatin-expressing (SOM) neurons across layers 2/3, 4, and 5/6. Unknown parameters, including the connection strength between the populations, were fitted to the data. Our results revealed similar population dynamics, fitted model parameters, predicted equivalent current dipoles (ECD), tuning curves, and lateral inhibition profiles across recording sites and animals, in spite of quite different extracellular current distributions. We found that PV firing rates were higher in BF than in non-BF responses, mainly due to different strengths of tonotopic thalamic input, whereas SOM firing rates were higher in non-BF than in BF responses due to lateral inhibition.

In conclusion, we demonstrate the feasibility of the model-fitting approach in identifying the contributions of cell-type specific population activity to stimulus-evoked LFPs across cortical laminae, providing a foundation for further investigations into the dynamics of neural circuits underlying cortical sensory processing.

## 1 Introduction

Neural responses to sensory stimuli have been extensively studied in order to elucidate how the brain represents features of the environment. Evoked responses are a specific kind of *event-related* signal that reflect (mostly electrical) brain activity in response to stimuli, such as tactile impulses, images, or sounds. In humans, cortical evoked responses can be recorded noninvasively via electroencephalography (EEG) and magnetoencephalography (MEG), which are widely used methodologies for probing brain function in health and disease. In order to draw valid and informative conclusions from these noninvasively recorded signals, it is important to understand the neural mechanisms underlying their generation. The cortical sources of EEG and MEG are thought to be intracellular currents primarily associated with postsynaptic potentials in pyramidal neurons (Lopes da Silva, 2013; Næss et al., 2021; Vaughan H.G.Jr, 1988). Pyramidal neurons and interneurons constitute multiple distinct populations in different layers of the cortex, which are locally and globally interconnected in a recurrent fashion. Already at the local level, these recurrent networks implement important functions (e.g., Chien et al., 2019; Hahn et al., 2022; Kunze et al., 2019). In order to eventually map the observed evoked responses onto these functions, it is crucial to obtain detailed information about how various neuronal cell types and synaptic connections contribute to their generation.

Invasive studies in animal models can contribute substantially to this aim by allowing the recording/imaging of layer-specific local field potentials (LFP) and cell-type specific neural firing (see Sections 1.1 and 1.2). Such animal model studies have already provided considerable information about lamina-specific neural activity (Bruyns-Haylett et al., 2017; Hajizadeh et al., 2019, 2021, 2022; Kohl et al., 2022; Neymotin et al., 2020; Sumner et al., 2021) and enhanced our understanding of the functional roles of various types of inhibitory interneurons in cortical processing (Aponte et al., 2021; Blackwell & Geffen, 2017; Liu et al., 2019; Liu & Kanold, 2021; Studer & Barkat, 2022). However, to more thoroughly elucidate the underlying neural generators of evoked responses and their relationship with information processing in the brain, it is necessary to mechanistically link together the available information about cell types and local neuronal circuits in the brain, intracranially-recorded LFPs, and extracranially-measured EEG and MEG signals. This task requires computational models. Although several efforts in this direction have been made (see Section 1.3), most modeling studies, such as spiking-based single-column models or rate-based multi-column models, are limited by being purely forward simulations without fitting the models to actual recordings of brain activity (e.g., LFPs and EEG/MEG), leaving their proposed theories less mechanistically and empirically grounded.

### 1.1 Layer-specific data - local field potentials

Linear-array multi-channel electrodes are a unique methodological tool which allows the simultaneous recording of LFPs across cortical layers (e.g., Fishman et al., 2001; Schroeder et al., 1998; Steinschneider et al., 2003). The high spatial resolution of LFPs provides valuable information regarding the intracranial generators of event-related potentials/fields (ERPs/ERFs) and information flow within and across cortical layers. Multi-unit activity (MUA), which can be extracted from high-frequency components of LFPs, reflects the spiking of local neuron populations in the vicinity of each electrode contact. Current source density (CSD), the second spatial derivative of the LFPs, provides information about the net transmembrane currents that give rise to the measured LFPs (Mitzdorf, 1985; Pitts, 1952). MUA and CSD provide complementary insights into the dynamics of activity within local neural circuits, as MUA primarily reflects suprathreshold neuronal firing (output), while CSD primarily reflects current flow associated with synaptic input, such as excitatory and inhibitory post-synaptic potentials (EPSPs/IPSPs). However, the disentanglement of meaningful functional components of MUA and CSD derived from trans-laminar LFP signals is an ill-posed problem. MUA contains spikes originating from different populations of excitatory and inhibitory neurons, and CSD is a spatial mixture of extracellular sinks and sources that can be either active (i.e., synaptic activity) or passive (i.e., return currents). Excitatory synaptic activity results in an active sink and a passive source, whereas inhibitory synaptic activity results in an active source and a passive sink. Moreover, the interpretation of activity (especially CSD) at later latencies is more uncertain due to the involvement of long-range cortical inputs (Happel et al., 2010). So far, evoked CSD and MUA have been extensively used to characterize tuning curves (Fishman et al., 2000a, 2000b; Fishman & Steinschneider, 2006, 2009; Steinschneider et al., 1998), differentiate responses to different stimuli (e.g., best-frequency [BF] vs. non-BF and standard vs. deviant) (Fishman & Steinschneider, 2012; Lakatos et al., 2020; O’Connell et al., 2011; Schaefer et al., 2015), and compare local neural population responses in different brain regions (e.g., core vs. belt regions of auditory cortex) (Banno et al., 2022). Statistical analyses in these studies mostly focused on neuronal activity occurring at specific latencies (usually within 50ms) in specific cortical layers (usually layer 4 or 2/3). An alternative approach, called laminar population analysis (LPA), was proposed to decompose the recorded LFP and MUA into firing rates of multiple neural populations and corresponding spatial profiles (Einevoll et al., 2007; Głąbska et al., 2014; Głąbska et al., 2016). However, the extracted components were only mapped to excitatory populations, and the connections between populations were indirectly estimated by a template-fitting analysis. In short, despite the high spatial specificity provided by LFPs and CSD analysis, our understanding of the information flow within neural circuits that give rise to these recorded signals is limited by the underdetermined inverse problem (Tenke & Kayser, 2012).

### 1.2 Cell type-specific activity in auditory cortex

Calcium imaging and optogenetic techniques make it possible to observe the activity of specific neuron types in auditory cortex. The activity of pyramidal cells is actively shaped by inhibitory interneurons (Liu et al., 2019). The major inhibitory interneuron types have specific characteristics with regard to morphology, targets, electrophysiology, and plasticity (Studer & Barkat, 2022), and appear to play distinct functional roles in cortical information processing (Blackwell & Geffen, 2017). Parvalbumin-expressing (PV) interneurons are fast-spiking neurons and mostly show higher spontaneous and tone-evoked firing rates than excitatory neurons. They target the soma, proximal dendrites, and initial segment of the axon of excitatory neurons, providing efficient and strong inhibition to excitatory neurons within the cortical column (a radius of up to 130 μm). PV interneurons receive inputs from both local excitatory neurons and the thalamus (MGBv). Functionally, PV interneurons are thought to contribute to the balance of excitation and inhibition and to the control of bottom-up (feedforward) information flow (Hamilton et al., 2013). Somatostatin-expressing (SOM) interneurons, on the other hand, show lower spontaneous and tone-evoked firing rates than PV interneurons. They target the distal dendrites of excitatory neurons, and their inhibition can reach widely (a radius of up to 300 μm) along the tonotopic axis. SOM interneurons receive excitatory inputs mainly within the cortex and much less from the thalamus (Ji et al., 2016). The synapses from excitatory neurons to SOM interneurons are short-term facilitating, whereas many other classes of synapses (e.g., the excitatory synapses between pyramidal cells and fast-spiking PV neurons) are short-term depressing (Hayut et al., 2011). SOM interneurons show slower activation dynamics and wider lateral inhibition than PV interneurons, which suggests a functional role of SOM neurons in integrating information over temporal and spectral domains (Yavorska & Wehr, 2016). PV and SOM interneurons mutually inhibit each other (Walker et al., 2016; Yavorska & Wehr, 2016). Both PV and SOM interneurons are inhibited by vasoactive-intestinal-peptide-expressing (VIP) interneurons which provide cross-modal modulation of sensory coding (Bigelow et al., 2019). The interaction among these various types of interneurons (with different time scales in spiking pattern and plasticity) can lead to complex dynamics and may implement specific neural computations. Modeling approaches provide a good way to identify the functional roles of each type of interneuron. For example, Park and Geffen built minimal rate-based and spiking models that incorporate a simple PV-SOM compensating mechanism to account for experimental findings in auditory studies, such as stimulus-specific adaptation, forward suppression, and tuning-curve adaptation (Park & Geffen, 2020). Hahn and colleagues used a rate-based mean-field modeling approach to study the interaction between pyramidal cells as well as SOM, PV, and VIP interneurons in different layers, which implements ultrasensitive switches toggling pyramidal neurons between high and low firing rate states associated with oscillations in distinct frequency bands (Hahn et al., 2022). Specifically, the pyramidal upstate is dominated by PV mediated high-frequency (gamma) activity, while the downstate is characterized by SOM mediated low-frequency (beta) oscillations.

### 1.3 Existing biological models

Biological models have been used to bridge the gap between microscopic properties (e.g., neuron types, single-unit neurophysiology, and morphology) and meso-/macroscopic observations (e.g., LFP/EEG/MEG signals). These models vary in their level of detail and scope. For example, a single-column model of the primary visual cortex (consisting of excitatory and inhibitory neurons in layers 2/3, 4, 5, and 6) was constructed to simulate laminar LFPs under different input conditions (Hagen et al., 2016, 2018). A single-column model of the primary auditory cortex (consisting of E, PV, SOM, VIP, and Neurogliaform cells in six layers) was recently constructed to simulate LFP, CSD, and EEG signals that replicate many experimental observations such as spontaneous neural activity and evoked responses to speech input (Dura-Bernal et al., 2022). Such detailed single-column models predict the contribution of layer- and cell-type-specific neuronal populations when the simulations match experimental observations. Some other, less detailed, models have been constructed to account for the generation of evoked responses. A single-column model (consisting of excitatory and inhibitory neurons in layers 2/3 and 5) using Human Neocortical Neurosolver suggests the contribution of a sequence of bottom-up thalamic inputs (targeting the soma of pyramidal neurons and causing upward currents) and top-down cortical inputs (targeting dendrites of pyramidal neurons and causing downward currents) (Kohl et al., 2022; Lakatos et al., 2020). This single-column model relates the ERP/ERF to intracellular currents in pyramidal long dendrites but leaves the origins of the sequences of inputs unexplained. This issue was addressed by a rate-based core-belt-parabelt model that includes an entire network of brain regions comprising auditory cortex (208 cortical columns, each column consisting of one excitatory and one inhibitory population), where ERFs are considered as the weighted sum of spatially distributed damped harmonic oscillators emerging out of coupled excitation and inhibition (Hajizadeh et al., 2019, 2021, 2022). This model provides a holistic perspective on the generation of ERFs. However, the proposed damped modes are extractions from the whole network dynamics, which can be difficult to link with LFP observations for validation. There remain several principal shortcomings in modeling evoked responses. On the one hand, models that take into account neuronal details (e.g., various types of inhibitory neurons) seemingly lack sufficient consideration of inter-column or inter-area interaction. On the other hand, models that focus on network dynamics and mechanisms currently provide less information about the role of different inhibitory neurons, and the simulations only qualitatively match the LFP/EEG/MEG recordings. Hence, there is a need for biological models that incorporate both a high degree of detail and a broad scope to clarify the neural underpinnings of cortical evoked responses.

### 1.4 Goals and approach of this study

In this study, we attempt to overcome the above-mentioned limitations by developing a multi-column model of auditory cortex with sufficient detail regarding neural populations and their interconnections to quantitatively reproduce layer-specific intracranial LFP recordings and qualitatively explain extracranially observable evoked responses. We utilized a biological cortical column model accounting for cell-type-specific interactions, which was integrated into a minimalistic multi-column array, representing the most relevant aspects of cortical architecture with respect to the tonotopic processing of auditory stimuli. Model parameters were specified by fitting the model to LFPs of tone-evoked responses simultaneously recorded across the layers of primary auditory cortex (A1) of awake monkeys. We show that the proposed model not only consistently replicates and explains detailed features of tone-evoked LFPs in A1, but also reproduces relevant aspects of extracranially-recorded evoked responses and predicts cell-type specific contributions to these signals.

## 2 Methods

### 2.1 Experimental data

#### 2.1.1 Acquisition and preprocessing

Neurophysiological data were obtained from A1 in 3 adult male macaque monkeys (Macaca fascicularis) using previously described methods (Fishman & Steinschneider, 2010; Steinschneider et al., 2003). All experimental procedures were reviewed and approved by the Association for Assessment and Accreditation of Laboratory Animal Care-accredited Animal Institute of Albert Einstein College of Medicine and were conducted in accordance with institutional and federal guidelines governing the experimental use of non-human primates. Animals were housed in our Association for Assessment and Accreditation of Laboratory Animal Care-accredited Animal Institute under daily supervision of laboratory and veterinary staff. Before surgery, monkeys were acclimated to the recording environment and were trained to perform a simple auditory discrimination task (see below) while sitting in custom-fitted primate chairs.

##### Surgical procedure

Under pentobarbital anesthesia and using aseptic techniques, rectangular holes were drilled bilaterally into the dorsal skull to accommodate epidurally placed matrices composed of 18-gauge stainless steel tubes glued together in parallel. Tubes served to guide electrodes toward A1 for repeated intracortical recordings. Matrices were stereotaxically positioned to target A1 and were oriented to direct electrode penetrations perpendicular to the superior surface of the superior temporal gyrus, thereby satisfying one of the major technical requirements of one-dimensional current source density (CSD) analysis (Müller-Preuss & Mitzdorf, 1984; Steinschneider et al., 1992). Matrices and Plexiglas bars, used for painless head fixation during the recordings, were embedded in a pedestal of dental acrylic secured to the skull with inverted bone screws. Perioperative and postoperative antibiotic and anti-inflammatory medications were always administered. Recordings began after at least 2 weeks of postoperative recovery.

##### Stimuli

Stimuli were generated and delivered at a sample rate of 48.8 kHz by a PC-based system using an RX8 module (Tucker Davis Technologies). Frequency response functions (FRFs) derived from responses to pure tones characterized the spectral tuning of the cortical sites. Pure tones used to generate the FRFs ranged from 0.15 to 18.0 kHz, were 200 ms in duration (including 10 ms linear rise/fall ramps), and were randomly presented at 60 dB SPL with a stimulus onset-to-onset interval of 658 ms. Resolution of FRFs was 0.25 octaves or finer across the 0.15–18.0 kHz frequency range tested. All stimuli were presented via a free-field speaker (Microsatellite; Gallo) located 60 degrees off the midline in the field contralateral to the recorded hemisphere and 1 m away from the animal’s head (Crist Instruments). Sound intensity was measured with a sound level meter (type 2236; Bruel and Kjaer) positioned at the location of the animal’s ears. The frequency response of the speaker was flat (within 5 dB SPL) over the frequency range tested.

##### Recordings

Neurophysiological recordings were conducted in an electrically shielded, sound-attenuated chamber. Monkeys were monitored via video camera throughout each recording session. To promote alertness and attention to the sounds during the recordings, animals performed a simple auditory discrimination task (detection of a randomly presented noise burst interspersed among test stimuli) to obtain liquid rewards.

Local field potentials (LFPs) and multiunit activity (MUA) were recorded using linear-array multi-contact electrodes, comprising 16 contacts, evenly spaced at 150-micron intervals (U-Probe; Plexon). Individual contacts were maintained at an impedance of 200 kΩ. An epidural stainless-steel screw placed over the occipital cortex served as the reference electrode. Neural signals were bandpass filtered from 3 Hz to 3 kHz (roll-off 48 dB/octave) and digitized at 12.2 kHz using an RA16 PA Medusa 16-channel preamplifier connected via fiber-optic cables to an RX5 data acquisition system (Tucker-Davis Technologies). LFPs time-locked to the onset of the sounds were averaged on-line by computer to yield auditory evoked potentials (AEPs). CSD analyses of the AEPs characterized the laminar distribution of net current sources and sinks within A1 and were used to identify the laminar location of concurrently recorded AEPs and MUA (Steinschneider et al., 1992, 1994). CSD was calculated using a 3-point algorithm that approximates the second spatial derivative of voltage recorded at each recording contact (Freeman & Nicholson, 1975; Nicholson & Freeman, 1975). MUA used to characterize the frequency tuning of each recording site (i.e., electrode penetration) was derived from the spiking activity of neural ensembles recorded within lower lamina 3, as identified by the presence of a large-amplitude initial current sink that is balanced by concurrent superficial sources in mid-upper lamina 3 (Fishman et al., 2001; Steinschneider et al., 1992). This current dipole configuration is consistent with the synchronous activation of pyramidal neurons with cell bodies and basal dendrites in lower lamina 3. Previous studies have localized the initial sink to the thalamorecipient zone layers of A1 (Metherate & Cruikshank, 1999; Müller-Preuss & Mitzdorf, 1984; Steinschneider et al., 1992; Sukov & Barth, 1998). To derive MUA, neural signals (3 Hz to 3 kHz pass-band) were high-pass filtered at 500 Hz (roll-off 48 dB/octave), full-wave rectified, and then low-pass filtered at 520 Hz (roll-off 48 dB/ octave) before averaging of single-trial responses (for a methodological review, see Supèr & Roelfsema, 2005). MUA (referred to as MUA_E in Supèr & Roelfsema, 2005) is a measure of the envelope of summed (synchronized) action potential activity of local neuronal ensembles (Brosch et al., 1997; O’Connell et al., 2011; Schroeder et al., 1998; Supèr & Roelfsema, 2005; Vaughan H.G.Jr, 1988). Thus, whereas firing rate measures are typically based on threshold crossings of neural spikes, MUA, as derived here, is an analog measure of spiking activity in units of response amplitude (e.g., see Kayser et al., 2007). MUA and single-unit activity, recorded using electrodes with an impedance similar to that in the present study, display similar orientation and frequency tuning in primary visual and auditory cortex, respectively (Kayser et al., 2007; Supèr & Roelfsema, 2005). Adjacent neurons in A1 (i.e., within the sphere of recording for MUA) display synchronized responses with similar spectral tuning, a fundamental feature of local processing that may promote high-fidelity transmission of stimulus information to subsequent cortical areas (Atencio & Schreiner, 2013).

Positioning of electrodes was guided by online examination of click-evoked AEPs. Pure tone stimuli were delivered when the electrode channels bracketed the inversion of early AEP components and when the largest MUA and initial current sink were situated in middle channels. Evoked responses to 40 presentations of each pure tone stimulus were averaged with an analysis time of 500 ms that included a 100 ms pre-stimulus baseline interval. The BF of each cortical site was defined as the pure tone frequency eliciting the maximal MUA within a time window of 0-75 ms after stimulus onset. As it was very consistent across layers (see Fig. S1), we used, for each recording site, the electrode channel with the maximum MUA for BF determination. The response time window includes the transient “On” response elicited by sound onset and the decay to a plateau of sustained activity in A1 (e.g., see (Fishman & Steinschneider, 2009)).

The neural responses to pure tones were acquired at the beginning of each recording session, after which the animals participated in at least three different auditory experiments. Finally, the monkeys were deeply anesthetized with sodium pentobarbital and transcardially perfused with 10% buffered formalin. Tissue was sectioned in the coronal plane (80-µm thickness) and stained for Nissl substance to reconstruct the (Morel et al., 1993) electrode tracks and to identify A1 according to previously published physiological and histological criteria (Kaas & Hackett, 2000; Merzenich & Brugge, 1973; Morel et al., 1993). Based upon these criteria, all electrode penetrations considered in this report were localized to A1, although the possibility that some sites situated near the low-frequency border of A1 were located in field R cannot be excluded.

#### 2.1.2 Preparation for model fitting

The target data for model fitting are comprised of the laminar MUA and CSD profiles of evoked responses to the best frequency (BF, see an example in Fig. 1) and 4 non-BF tones at four recording sites (site 1 from monkey A, site 2 from monkey D, sites 3 and 4 from monkey E). The 4 sites were selected because they display stereotypical features of laminar AEP, MUA, and CSD neurophysiological response profiles which are characteristic of primary auditory cortex (A1), as described in previous publications by various investigators (Fishman, 2014; Fishman et al., 2000b; O’connell et al., 2014). Moreover, the different BFs of these sites lie within the frequency range of maximum hearing sensitivity in macaques. Hence, we regard these sites as representative of population responses in A1, which are well-suited for the computational modeling of auditory cortical responses. See Fig. S2 for further details. For each recording site, the experimental data (BF and 4 non-BF responses) were cropped (time window: 0 to 200 ms, 200 timepoints), concatenated along the time axis and normalized to their maximum peak, resulting in a target MUA matrix [1000 timepoints × 16 channels] and a target CSD matrix [1000 timepoints × 12 channels] for model fitting.

**Figure 1.**
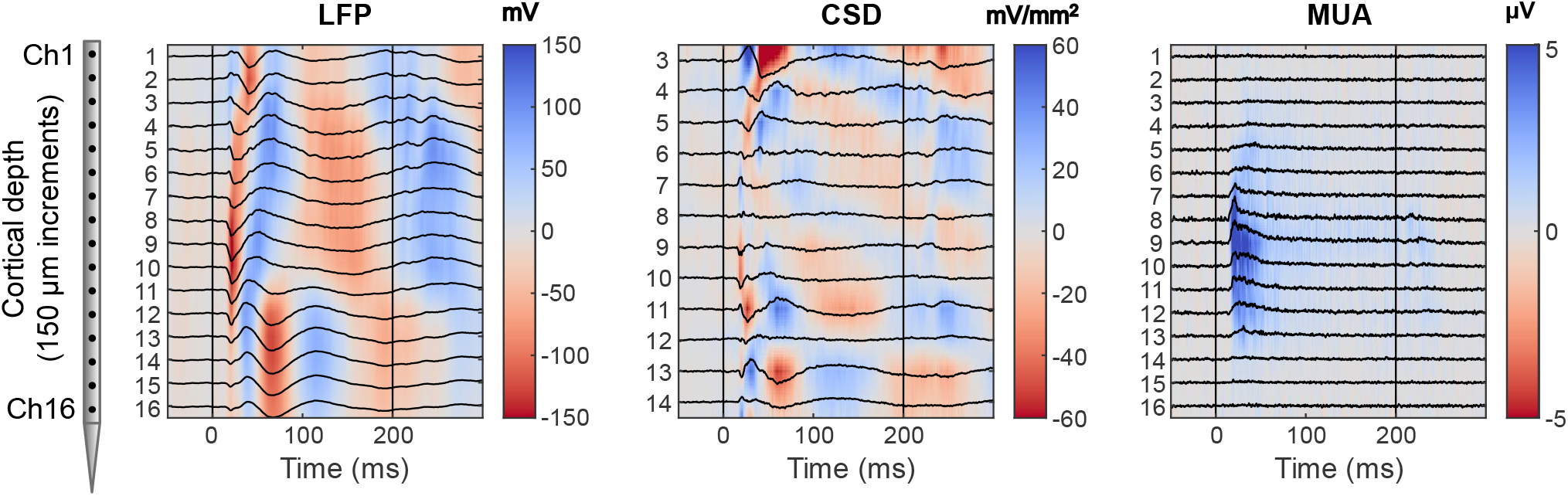
Laminar profiles of evoked responses to a BF tone recorded in a representative electrode penetration into A1. The 16-channel electrode array records local field potentials (LFPs) from superficial to deep layers (2000 µm), from which current source density (CSD) and multi-unit activity (MUA) are derived. A schematic of the electrode array is shown on the left of the figure. For visualization, the values of response amplitude are also color-coded (positive values in blue; negative values in red). Tone onset and offset times are indicated by the vertical drop lines.

### 2.2 Cortical column model

The cortical model consists of two columns. Each column contains 7 neural populations identified by cell type and laminar location of their cell bodies: excitatory pyramidal neurons (L2/3 E, L4 E, and L5/6 E), inhibitory PV interneurons (L2/3/4 PV and L5/6 PV), and inhibitory SOM interneurons (L2/3/4 SOM and L5/6 SOM). Each neural population is described by a rate-to-potential operator (Section 2.2.4) and a potential-to-rate operator (Section 2.2.5) as in the Jansen-Rit model (Jansen & Rit, 1995). Different types of inhibitory interneurons were considered because of their distinct characteristics regarding morphology, connection motif, target neurons/locations, plasticity, spiking rate, synaptic time constant, and afferent inputs (Studer & Barkat, 2022), making it likely that they differentially contribute to the observed MUA and CSD. Thus, the model was designed with the aim of reconstructing the dynamics of different interneuron types from the LFP data. To reduce model complexity, VIP interneurons were not included, as VIP interneurons are thought to be involved in cortico-cortical interaction and neuromodulatory control (Mesik et al., 2015). We would like to note that, within our model framework, the number of neural populations is flexible, and its modification would only lead to different degrees of granularity in the spatiotemporal decomposition of MUA and CSD. In the model, we merge neurons into L2/3, L4, and L5/6 to characterize the dynamics at the spatial resolution of supragranular, granular, and infragranular layers.

As illustrated in Fig. 2A, column 1 represents the recording site, which does not necessarily respond optimally to the presented frequency (non-BF site), while column 2 (BF site) represents a location (cortical column) that responds optimally to the presented tone stimulus. For simplicity, the configurations of the two columns were set to be identical, and the two columns inhibit each other through symmetric inter-column E-to-SOM connections. The E and PV populations receive direct thalamic input with a fixed ratio based on the literature (see Section 2.2.2). Column 2 (BF site) receives the full strength of thalamic input (in=1), and column 1 (recording site) receives weaker input (in≤1), based on the tonotopic organization of the auditory pathway. The two-column model is a simplification of the anatomical and functional organization of A1, where the interactions with other cortical areas (e.g., non-primary fields of auditory cortex) are not considered. In the case of the BF response, both columns receive the same input strength (in=1), as they represent the same location and inter-column connections are in fact intra-column connections. This way, the same model could be used for both BF and non-BF situations.

**Figure 2.**
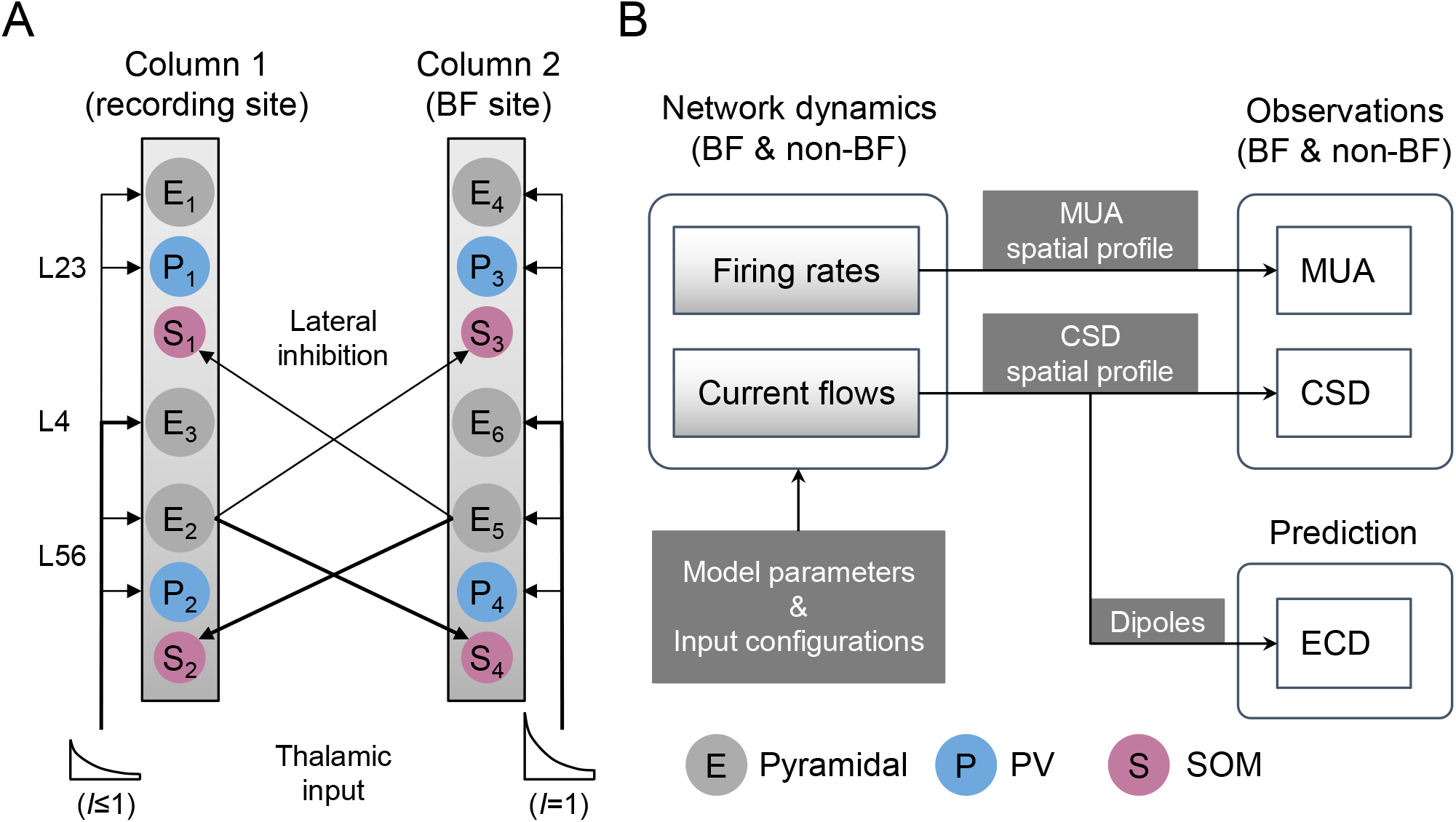
Two-column cortical model and forward simulation. (A) The model consists of two identical columns, where Column 1 represents the cortical area at the recording site, and Column 2 represents the latent cortical area that responds most strongly to the presented tonal stimulus (at the BF). The thalamic inputs make synaptic connections with the E and PV populations at L2/3, L4, and L5/6. The lateral inhibition consists of the connections from L5/6 E to the SOM populations at L2/3 and L5/6. For clarity, intra-column connections are not shown. Different line widths and circle sizes indicate the difference in connection strength and contribution to MUA, respectively (details in Tables 1 and 5). (B) The forward simulation includes two stages. First, the network dynamics are simulated based on the model parameters and input configurations under BF and non-BF conditions at the recording site. Second, the observation model translates firing rates and current flows to the observations (i.e., MUA and CSD) through the spatial profiles calculated by constrained least square fit. The current flows and CSD can be further translated to dipole signals as model predictions.

The model simulates the time series of latent variables including firing rates, postsynaptic potentials (PSP), and connectivity efficacy (related to short-term plasticity) under BF and non-BF conditions given the same model parameter settings but different input configurations. As illustrated in Fig. 2B, the network dynamics are transformed into simulated MUA and CSD via a forward model (see Section 2.3) to match the target data. The forward model also transforms network dynamics to a simulated equivalent current dipole (ECD) signal, which is equivalent (up to a scaling factor determined by the volume conduction of the head) to EEG/MEG signals. Thus, the model could be used to identify the specific contributions of different cell types to those signals.

#### 2.2.1 Thalamic input

The thalamic input *i*_*th*_(*t*) to the two-column model is a decay function with a delay (Eq. (1)). The parameters include thalamic input strength *I*, decay time constant *τ*_*in*_, decay level α, and delay *t*_*d*_. This is to mimic the fast decaying firing rate observed in the auditory system (Pérez-González & Malmierca, 2014).

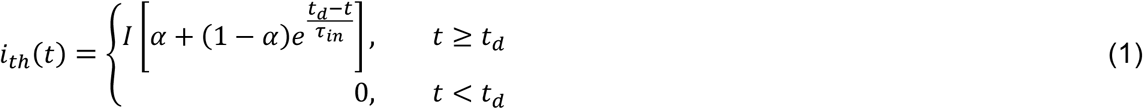

#### 2.2.2 Connectivity

The intra-column connectivity is derived from the reported connection probabilities and synaptic strengths in the primary visual cortex of mice (Billeh et al., 2020). The calculation is illustrated in Fig. 3, while the exact connectivity values can be found in the supplementary Table S1. The connectivity is grouped into default connectivity matrices *W*_*EE*_, *W*_*PE*_, *W*_*SE*_, *W*_*EP*_, *W*_*PP*_, *W*_*SP*_, *W*_*ES*_, and *W*_*PS*_, which can then be separately rescaled by free parameters *θ*_1−8_ in the optimization. For example, the 2-by-matrix *W*_*PE*_ (connections from E1, E2, and E3 to P1 and P2) is rescaled by *θ*_2_. The inter-column connections only consist of E-to-SOM connections *W*_*inter*−*SE*_ (from E2 in column 1 to SOM1 and SOM2 in column 2, and vice versa), which can be rescaled by parameters *θ*_20−24_ that control the strength of lateral inhibition under BF and non-BF conditions. The scaling parameters also allow accounting for potential discrepancies between the intra- and inter-columnar connectivity between mice and primates. Including inter-column E-to-E connections *W*_*inter*−*EE*_ does not significantly improve model performance and the resulting parameters (as shown in supplementary Fig. S3 that tests variant models), so we decided to keep the model simple.

**Figure 3.**
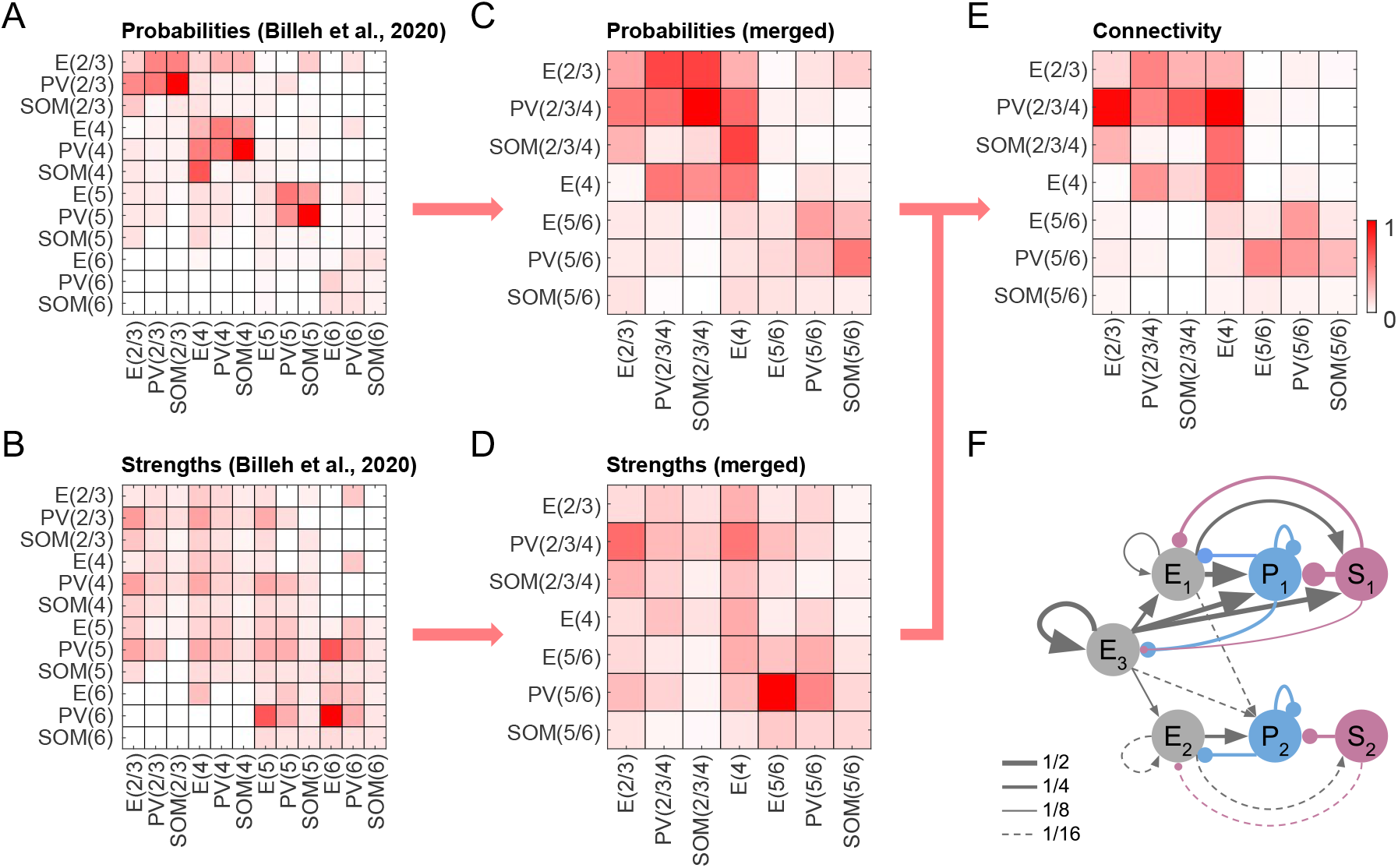
Default intra-column connectivity. (A, B) The connection probabilities and synaptic strengths are taken from (Billeh et al., 2020). The numbers in the parentheses stand for the cortical layers. (C, D) The probabilities and strengths are merged by averaging into coarser layers matching our model definitions. (E) The default intra-column connectivity is defined as the product of probabilities and strengths. Matrices A to E are normalized for visualization. (F) The default intra-column connectivity is visualized in a graph where the edge widths discretize the normalized connectivity values in matrix F. Values above 1/2 have the thickest edges, while values lower than 1/16 are not shown in the graph. For detailed values of connection probabilities, synaptic strengths, and connectivity, please refer to supplementary Table S1.

The thalamic input is fed only to E and PV populations. The input connectivity across layers (as listed in Table 1) is based on the normalized peak amplitude of thalamocortical responses and the laminar pattern of thalamocortical innervation in mouse primary auditory cortex reported in (Ji et al., 2016). The default thalamic input connectivity matrices *W*_*Eth*_ and *W*_*Pth*_ refer to connections from the thalamus to E1, E2, and E3, and to PV1 and PV2, respectively, which can be rescaled by parameters *θ*_9,10_ in the optimization.

**Table 1.** Thalamic input connectivity across laminar layers (normalized to E3).

| Neural population | (a) Normalized peak amplitude | (b) % Innervated | (a*b) Input connectivity |
| --- | --- | --- | --- |
| E1 (L2/3) | 129/418=0.3 | 75% | 0.225 |
| E3 (L4) | 418/418=1 | 100% | 1 |
| E2 (L5/6) | 163/418=0.4 | 85% | 0.34 |
| PV1 (L2/3/4) | 615/418=1.47 | 85% | 1.25 |
| PV2 (L5/6) | 426/418=1.02 | 100% | 1.02 |

#### 2.2.3 Rate-to-potential operator

The rate-to-potential operator implements the transformation of the firing rate *r*_*j*_(t) of the *j*th population to the postsynaptic potential *v*_*ij*_(t) at the *i*th population through an effective connection strength *w*_*ij*_(t). This transformation is achieved by convolving the weighted firing rate *w*_*ij*_(t)*r*_*j*_(t) with a synaptic kernel *h*_*ij*_(*t*) (Eq. (2), as in Chien et al., 2019; Jansen & Rit, 1995; Wang & Knösche, 2013). Note that the effective connection strength *w*_*ij*_(*t*) can become a variable, if short-term plasticity is taken into account (see Section 2.2.5).

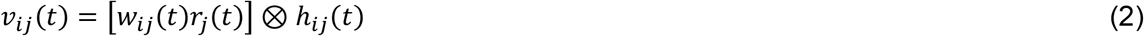

The synaptic kernel *h*(t) is described as a bi-exponential function parameterized by the scaling factor *H* and the time constants *τ*_1_ and *τ*_2_ (Eq. (3)).

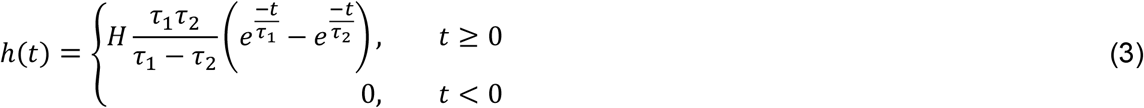

The convolution in Eq. (2) can be numerically realized by two first-order ordinary differential equations (Eqs. (4) and (5)).

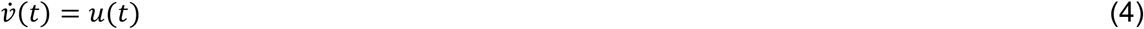

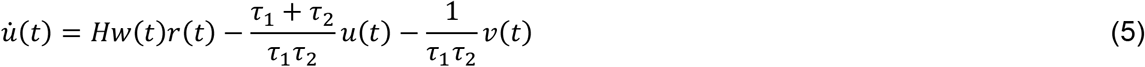

The shape of synaptic kernels depends on pre- and postsynaptic properties of the neurotransmitter-receptor systems, and the respective parameters *H*, *τ*_1_, and *τ*_2_ are listed in Table 2.

**Table 2.** Parameters of the synaptic-dendritic kernels for different connections.

| Synaptic type | | $H$ | $\tau_1(\text{ms})$ | $\tau_2(\text{ms})$ | Reference |
| --- | --- | --- | --- | --- | --- |
| $E \rightarrow E^{(1)}$ | AMPA | 14400 | 1 | 5.3 | (Dura-Bernal et al., 2022) |
|  | NMDA | 1200 | 3 | 70 |  |
| $E \rightarrow PV$ | NMDA | 7250 | 2.1 | 5.6 | (Jouhanneau et al., 2018) |
| $E \rightarrow SOM$ | NMDA | 3090 | 4.5 | 25.2 | (Jouhanneau et al., 2018) |
| $PV \rightarrow E$ | GABA<sub>A</sub> | -4000 | 1 | 18.2 | (Dura-Bernal et al., 2022) |
| $PV \rightarrow PV$ | GABA<sub>A</sub> | -5530 | 3.5 | 5.5 | (Bacci et al., 2003) |
| $PV \rightarrow SOM$ | GABA<sub>A</sub> | -7380 | 1.4 | 101 | (Bacci et al., 2003) |
| $SOM \rightarrow E^{(2)}$ | GABA<sub>A</sub> | -1800 | 2 | 100 | (Dura-Bernal et al., 2022) |
|  | GABA<sub>B</sub> | -100 | 25 | 300 |  |
| $SOM \rightarrow PV$ | GABA<sub>A</sub> | -1800 | 2 | 100 | (Dura-Bernal et al., 2022) |

(1) $AMPA:NMDA=83:17$
(2) $GABA_A:GABA_B=50:50$

The PSPs *v*(*t*) are then used to calculate the current flows *c*(*t*). In Eq. (6), a current flow *c*_*j*_(*t*) caused by the *j*th input source (from a cortical or a thalamic input) is calculated as the sum of the absolute values of PSPs *v*_*ij*_(*t*) at all E populations (*N*_*E*_ = 3) at the recording site (i.e., Column 1). The mapping to MUA and CSD is illustrated in Fig. 4.

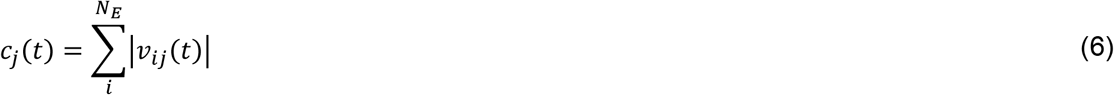

**Figure 4.**
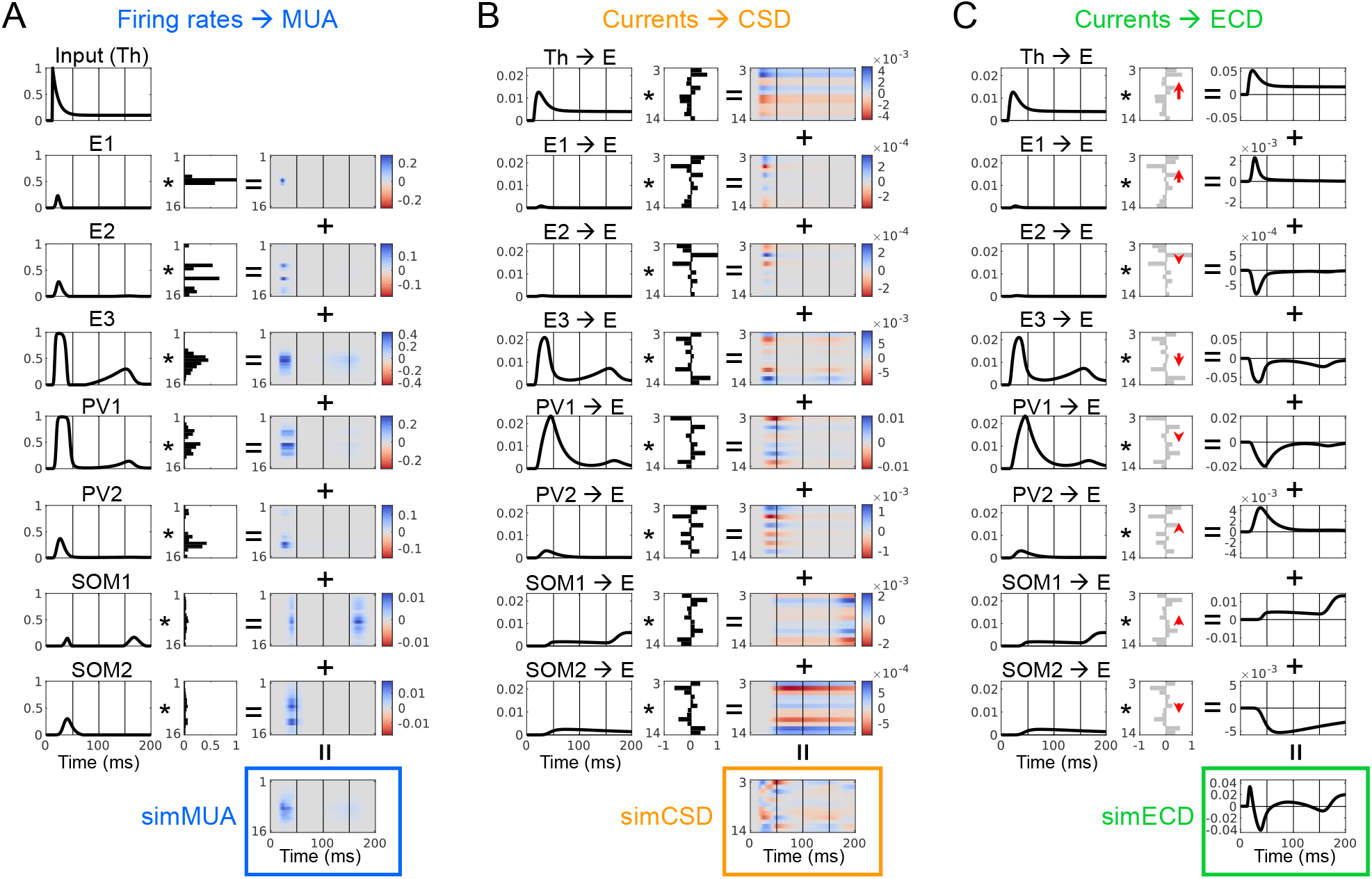
Forward mapping from network dynamics to observations. (A) The simulated MUA (simMUA) is the sum of individual firing rates (left column) weighted by the non-negative MUA spatial profile (middle column) of the recording site (Column 1). (B) The simulated CSD (simCSD) is the sum of individual current flows (left column) weighted by the CSD spatial profile (middle column) of the recording site (Column 1). (C) The simulated equivalent current dipole (simECD) is the sum of individual current flows (left column) weighted by dipole directions and lengths (red arrows in the middle column) derived from the CSD spatial profile (gray color in the middle column).

#### 2.2.4 Potential-to-rate operator

The potential-to-rate operator transforms the overall PSP *v*_*i*_(t) at the *i*th population into mean firing rate *r*_*i*_(*t*) using a sigmoid function (Eqs. (7) and (8)). The overall PSP is the sum of EPSPs and IPSPs caused by *N*_*cur*_ presynaptic current sources which include *N*_*pop*_ excitatory/inhibitory populations and thalamic input.

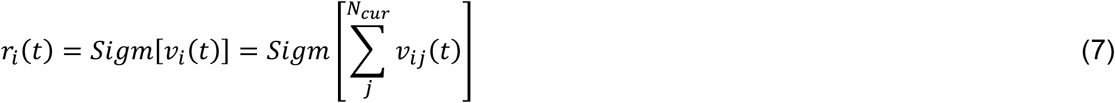

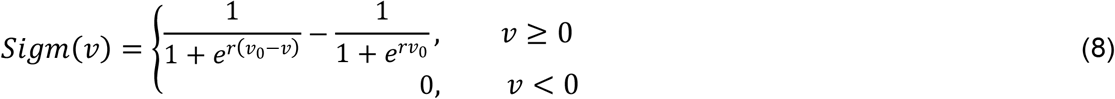

The output firing rate is limited to 0 ≤ *r*(*t*) ≤ 1, where *r* = 1 refers to the max firing rate of the E/PV/SOM neuron type. The sigmoid function is shifted so that *Sigm*(0) = 0, meaning no change in the firing rate relative to baseline. To make the fitting process more stable, negative firing rates are set to 0. The sigmoid functions for the E, PV, and SOM populations have different slopes *r* and thresholds *v*_0_ (as listed in Table 3) based on the firing characteristics of the neuron types reported in (Fanselow et al., 2008).

**Table 3.**
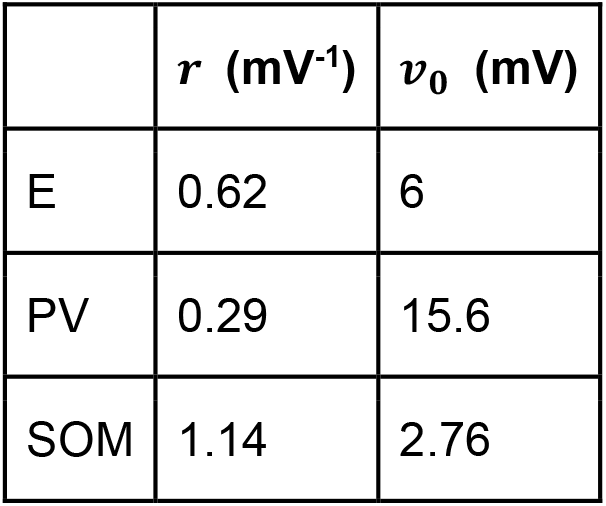
Parameters of the sigmoid functions.

#### 2.2.5 Short-term plasticity

Short-term plasticity (STP) refers to the modulation of synaptic strength based on the history of presynaptic activity. STP can be depressing (STD) or facilitating (STF), which depends on the synaptic type (Blackman et al., 2013; Fino et al., 2013; Ma et al., 2012; Regehr, 2012; Silberberg et al., 2005). STD can be modeled by a synaptic efficacy variable *x* (0 ≤ *x* ≤ 1), which denotes the fraction of remaining neurotransmitters (or synaptic vesicles). STF can be modeled by another synaptic utilization variable *u* (0 ≤ *u* ≤ 1) which denotes the neurotransmitters ready for use (release probability) (Silberberg & Markram, 2007). The two variables then modulate the fixed connection strength *w*_0_ as in Eq. (9).

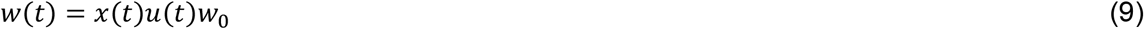

The variables *u*(*t*) and *x*(*t*) depend on the presynaptic firing rate *r*(*t*) as in Eqs. (10) and (11). The parameter *U* is the initial release probability, and parameters *τ*_*f*_ and *τ*_*d*_ denote the decay and recovery time constants, respectively. The parameters *k*_*f*_ and *k*_*d*_ are the change rates and are tuned in the optimization procedure.

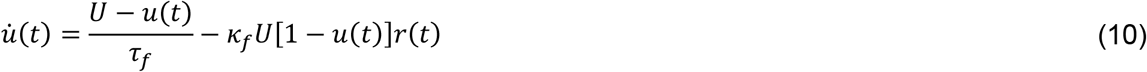

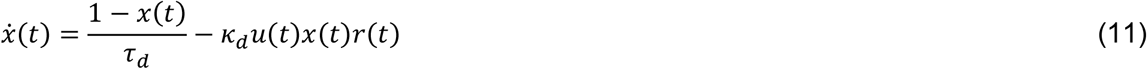

The effect of STP on the dynamics of a network of E, PV, and SOM populations has been studied by Hayut and colleagues (Hayut et al., 2011). Among the eight synaptic types, only E-to-SOM connections exhibit STF, while the remaining connections exhibit STD. To simplify the model, we only consider STF on E-to-SOM connections and STD on E-to-E connections in the model. These are two most prominent presynaptic plasticities in the neocortex reported in (Kalisman et al., 2005; Silberberg et al., 2005; Silberberg & Markram, 2007). The parameters of STP are based on (Hayut et al., 2011; Silberberg & Markram, 2007). See Table 4 for details.

**Table 4.** Parameters of short-term plasticity.

| | $U$ | $\tau_f$ (ms) | $\tau_d$ (ms) | $\kappa_f$ | $\kappa_d$ |
| --- | --- | --- | --- | --- | --- |
| E→E | 1 | - | 200 | 0 | 20* |
| E→SOM | 0.05 | 670 | - | 600* | 0 |

*Note: Values marked by \* are default values that will be rescaled by free parameters.*

### 2.3 Observation model

The observation model includes two spatial profiles which map the network dynamics to M A and CS. o match the target data, the network dynamics are subsampled to *T* = 200 timepoints (i.e., firing rates *r* into *S*_*rate*_ [*N*_*pop*_×*T*], and current flows *c* into *S*_*current*_ [*N*_*cur*_×*T*], where *N*_*pop*_ = 7 populations, and *N*_*cur*_ = 8 input sources.

#### 2.3.1 Spatial profiles

The MUA spatial profile *A*_*MUA*_ [*N*_*ch*_×*N*_*pop*_] describes the sensitivity of channels (*N*_*ch*_ = 16) to the firing rates of neural populations (see Section 2.3.2), which is closely related to the spatial distribution of cell bodies (or axonal hillocks, where the spikes are generated). The simulated MUA *Φ*_*MUA*_ [*N*_*ch*_×*T*] is the multiplication of the mixing matrix *A*_*MUA*_ and the firing rates *S*_*rate*_ (Eq. (12)).

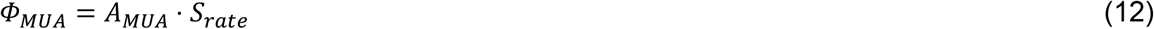

The CSD spatial profile *A*_*CSD*_ [*N*_*ch*′_×*N*_*cur*_] describes the sensitivity of the spatial distribution of sinks and sources along the channels (*N*_*ch*′_ = 12) to the current flows. The simulated CSD *Φ*_*CSD*_ [*N*_*ch*′_×*T*] is the multiplication of the mixing matrix *A*_*CSD*_ and the current flows *S*_*current*_ (Eq. (13)).

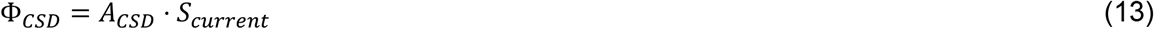

#### 2.3.2 Constraints on the spatial profiles

Both spatial profiles *A*_*MUA*_ and *A*_*CSD*_ are difficult to determine by measurements. They are estimated by constrained regression of the network dynamics *S*_*rate*_ and *S*_*current*_ to the target data 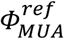 and 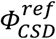. The goal is to find spatial profiles that minimize the distance between simulation *Φ* and target *Φ*^*ref*^, while, at the same time, obeying certain plausibility constraints.

The constraint on the MUA spatial profile *A*_*MUA*_ [*N*_*ch*_×*N*_*pop*_] is based on the fact that the sensitivity to the firing rate of a certain cell type is proportional to the product of cell density and maximal firing rate. Therefore, the ratio of the column sums of *A*_*MUA*_ should be fixed (Table 5). This is a relatively coarse constraint where the laminar distribution of cell density is not considered.

**Table 5.** Constraint of MUA spatial profile.

| Neural population | (a) Proportion of neurons<br>(Keller et al., 2018; Rudy et al., 2011) | (b) Maximal firing rate (Beierlein et al., 2003) | (a*b) Contribution to MUA<br>(normalized to E1) |
| --- | --- | --- | --- |
| E1, E2, and E3 | 1/3 of 80% | 59.4 Hz | 1 |
| PV1 and PV2 | 1/2 of 20%*40% | 271.7 Hz | 0.69 |
| SOM1 and SOM2 | 1/2 of 20%*30% | 120.7 Hz | 0.23 |

The constraint on the CSD spatial profile *A*_*CSD*_ [*N*_*ch*′_× *N*_*cur*_] is based on the fact that the transmembrane currents should be conserved. This means, the sources and the sinks need to match. Therefore, each column of *A*_*CSD*_ should add up to zero (i.e., area of sinks = area of sources). Additionally, in order to avoid ambiguity with the overall current magnitudes, the norms of all column vectors of *A*_*CSD*_ are constrained to be the same.

#### 2.3.3 Equivalent current dipole

The neuronal generators of EEG/MEG signals are commonly estimated as equivalent current dipoles (ECDs) by source localization techniques such as dipole fitting and beamforming. The simulated ECDs provide a theoretical link between LFPs and EEG/MEG signals, by which we can also examine the contribution of E, PV, and SOM populations to event-related deflections, like P1, N1, and N2.

The model predicts *Φ*_*ECD*_ [1×*T*] at the recording site once current flows *S*_*current*_ and a CSD spatial profile *A*_*CSD*_ are given. As in Eq. (14), *Φ*_*ECD*_ is calculated as the multiplication of 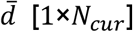 and the current flows *S*_*current*_ [*N*_*cur*_×*T*], where 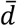 is the displacement from the mean center of sinks 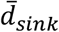 to the mean center of sources 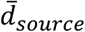 derived from each column of *A*_*CSD*_ [*N*_*ch*′_×*N*_*cur*_].

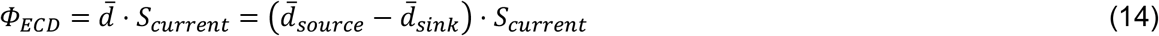

### 2.4 Optimization procedure

In general, we use a genetic searching scheme (Eiben & Smith, 2015; García-Martínez et al., 2018; Katoch et al., 2021) combined with a gradient descent (Gauss-Newton) method to optimize the 28 free parameters *θ* within a predefined search range (Table 6), in order to minimize the cost function *f*(*θ*) for each recording site. Since the cost function has a complicated shape with multiple local minima, we made use of the assumption that the true solutions should yield similar parameters across recording sites. In order to reliably find these solutions, we iteratively continued the search for each recording site starting at the solutions for all other recording sites. If our assumption is correct, this should lead to better fits for the individual recording sites.

**Table 6.** Free parameters and the corresponding default model parameters.

| Model parameters | Default values of model parameters | Free parameters | Ranges of free parameters [min, max] |
| --- | --- | --- | --- |
| Connectivity matrices $W_{EE}$, $W_{PE}$, $W_{SE}$, $W_{EP}$, $W_{PP}$, $W_{SP}$, $W_{ES}$, and $W_{PS}$ | Table. S1C | $\theta_{1-8}$ | [0.1, 15] |
| Thalamic input connectivity matrices $W_{Eth}$ and $W_{Pth}$ | Table 1 | $\theta_{9,10}$ | [0.1, 15] |
| Short-term plasticity $\kappa_{d,EE}$ and $\kappa_{f,SE}$ | Table 4 | $\theta_{11,12}$ | [0.5, 4] |
| Synaptic kernel time constants $\tau$ | Table 2 | $\theta_{13}$ | [0.5, 1.5] |
| Sigmoid function slopes $r$ | Table 3 | $\theta_{14}$ | [0.5, 1.5] |
| Thalamic input decay levels $\alpha_{1-5}$ (BF and non-BF conditions) | Equal to 1 | $\theta_{15-19}$ | [0.1, 0.5] |
| lateral inhibition $W_{inter-SE}$ for (BF and non-BF conditions) | Table. S1C | $\theta_{20-24}$ | [0.1, 15] |
| Thalamic input strengths $I_{1-4}$ to Column 1 (non-BF conditions) | Equal to 1 | $\theta_{25-28}$ | [0.1, 1] |

Note that this additional similarity criterion is only used to guide the search for the solutions, while the sole criteria for the goodness of fit are the cost functions for the individual recording sites. The optimization scheme is detailed in Fig. 5.

**Figure 5.**
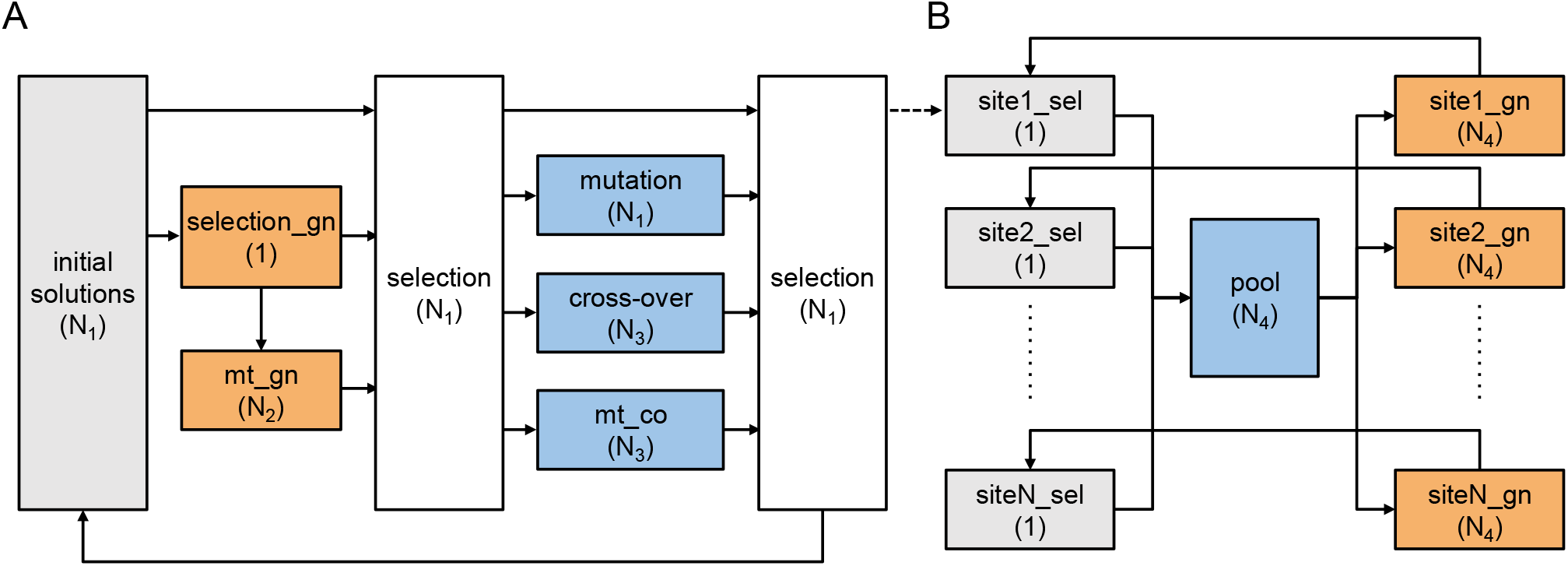
Flow chart of optimization procedure. (A) Optimization for an individual recording site. The iteration starts with N_1_ = 60 initial random solutions (parameter sets). The orange box ‘selection_gn’ selects the best solution (with highest R^2^) and fine-tunes that solution using the Gauss-Newton (gn) method to the next local minimum. The orange box ‘mt_gn’ mutates that locally optimal solution, one parameter at a time, resulting in N_2_ = 28 modified solutions. These are then fine-tuned again using the gn method. The white box ‘selection’ keeps the first N_1_ solutions with highest R^2^ from the N_1_ + 1 + N_2_ solutions. The blue box ‘mutation’ creates another N_1_ solutions, while ‘cross-over’ and ‘mt_co (mutated_cross-over)’ boxes each generate N_3_ random solutions (N_3_ = 2000). The next white box ‘selection’ again keeps the first (best) N_1_ solutions from these 2N_1_ + 2N_3_ solutions and replaces the initial solutions. The best solution for an individual site is selected after 10 iterations. (B) Optimization across recording sites. The iteration starts with the best solution for each recording site. Noise is added to these solutions with a uniform distribution (range: ±std(i), i ∈ 1,2, …, 28) to generate N_4_ = 1000 jittered solutions in the blue box ‘pool’. Each of these jittered solutions is then fine-tuned for each recording site with respect to R^2^ using the gn method (the orange boxes). The optimized solution for each recording site is updated iteratively if a fine-tuned solution with higher R^2^ is found. The iteration ends when there is no update. Then, for each recording site, the best-fitting solution is selected as the final solution. The number in the parentheses in each box indicates the resulting number of solutions.

The cost function *f*(*θ*) is defined as the sum of squared errors (SSE) between the simulation *Φ*(*θ*) and target *Φ*^*ref*^ (Eq. (15)). The target data *Φ*^*ref*^ comprises 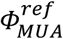 and 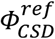 under BF and non-BF conditions. The benefit of including non-BF conditions in the optimization procedure is two-fold. First, the amount of target data is increased while the degrees of freedom of spatial profiles remain the same, which reduces the risk of overfitting. Second, the dynamics of PV and SOM populations under BF vs. non-BF conditions can be compared.

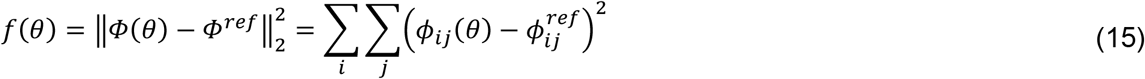

The simulated data *Φ*(*θ*) of an individual recording site comprises of *Φ*_*MUA*_(*θ*) and *Φ*_*CSD*_(*θ*) as in Eqs. (12) and (13). The forward simulation includes two stages. In the first stage, the default two-column model and input configuration are rescaled by 28 free parameters *θ* to generate network dynamics *S*_*rate*_ and *S*_*current*_. The 28 free parameters *θ* rescale intra-column connections *W*_*EE*_, *W*_*PE*_, *W*_*SE*_, *W*_*EP*_, *W*_*PP*_, *W*_*SP*_, *W*_*ES*_, and *W*_*PS*_ (*θ*_1−8_), thalamic input connectivity matrices *W*_*Eth*_ and *W*_*Pth*_ (*θ*_9,10_), rates of short-term plasticity *κ*_*d,EE*_ and *κ*_*f,SE*_ (*θ*_11,12_), synaptic time constants *τ* (*θ*_13_), slopes of sigmoid functions *r* (*θ*_14_), thalamic input decay levels α_1−5_ for BF and non-BF conditions (*θ*_15−19_), lateral inhibition *W*_*inter*−*SE*_ for BF and non-BF conditions (*θ*_20−24_), and thalamic input strengths to column 1 for non-BF conditions (*θ*_25−28_). We chose a reasonably larger range for connectivity-related free parameters (*θ*_1−10,20−24_) to allow enough flexibility of network dynamics. We chose a narrower range for the rest of the free parameters to adhere to the evidence from the literature. The search ranges of the 28 free parameters are summarized in Table 6. In the second stage, the spatial profiles *A*_*MUA*_ and *A*_*CSD*_ are optimized by regressing the resulting network dynamics *S*_*rate*_ and *S*_*current*_ to the target 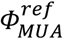 and 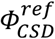, respectively. Multiplication of spatial profiles with network dynamics then yields the predicted MUA and CSD, which can be compared to the observed MUA and CSD.

We use the Gauss-Newton (gn) method to approach the minimum of the cost function *f*(*θ*) in Eq. (15). This method was successfully used in our previous studies for optimization of neural mass models with large numbers of free parameters (Wang et al., 2019; Wang & Knösche, 2013). The Jacobian matrix 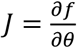 is numerically approximated with ewton’s difference quotient.

The genetic search scheme (see Fig. 5) includes three operating boxes for broadening the exploration in parameter space. In the ‘mutation’ box, the solutions are sorted based on the goodness-of-fit (cost). Then, each parameter of a solution can mutate (i.e., be replaced by a random value) with a chance that linearly ranges from 10% (for best fit case) to 90% (for worst fit case) based on the ranking of the solution.

In the ‘cross-over’ box, we randomly draw two solutions as parent solutions from the outputs of ‘selection’ and ‘mutation’ boxes. Then we randomly decide a cross-over point and generate two off-spring solutions by swapping the same sides of parameters between the parents (i.e., segment-wise swap). This is repeated to generate *N*_3_ solutions.

In the ‘mt_co’ box, we run the cross-over operator again. However, instead of swapping the whole right side of the cross-over point, we only swap the cross-over point (i.e., element-wise swap). This is repeated to generate *N*_3_ solutions. The parent solutions come from the output of ‘selection’, ‘mutation’, and ‘cross-over’ boxes.

The effect of model parameters on the network dynamics can be highly nonlinear, and a global minimum in cost function is hard to identify in high-dimensional space. In addition, a global minimum for a single recording site may still be at risk of overfitting. Therefore, it is important to check the similarity and stability of the common solutions. We explore the cost function surface by deviating each parameter at a time, starting from a final solution in the parameter space.

### 2.5 Non-negative matrix factorization

We examined whether a blind decomposition approach would yield similar predictions (e.g., ECD) as our model-fitting approach. The non-negative matrix factorization (a Matlab function *nnmf*) was used to decompose target 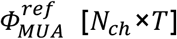 into two non-negative matrices 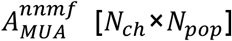 and 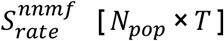 where the root mean square residual between 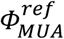 and 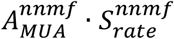 is minimized. The decomposed firing rates 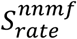 are then convolved with alpha kernels to generate current flows 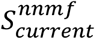. The time constants of the alpha kernels are optimized in similar procedure as in Fig. 5A to minimize the SSE between 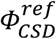 and 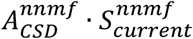. The ECD 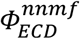 is then calculated in the same way as in Sections 2.3.3.

## 3 Results

We fitted the 28 parameters (see Methods section) of our cortical column model, such that the MUA and CSD derived from the electrophysiological data obtained at each recording site were explained best in a least squares sense. Concurrently with the parameters of the cortical column model, we also fitted a set of parameters of the observation model, namely the spatial sensitivity distribution of each neural population (i.e., MUA spatial profile *A*_*MUA*_) and the spatial distribution of current sinks and sources on pyramidal dendrites (i.e., CSD spatial profile *A*_*CSD*_). As a result, the fitted model predicted not only the MUA and CSD that we could compare to the empirically measured values, but also a set of latent (hidden) variables, namely the mean firing rates and the current flows of the neural populations.

In Section 3.1, we present the predicted data (*Φ*_*MUA*_ and *Φ*_*CSD*_) along with the target data (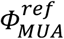and 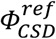) for a qualitative and quantitative comparison. In Section 3.2, we show the estimated latent variables, covering neural population activity (firing rates and current flows), as well as the ECD magnitude at the recording sites (ECD, *Φ*_*ECD*_). We also compare *Φ*_*ECD*_ with the ECD estimated by a non-negative decomposition approach 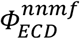 to evaluate the advantages of our model-fitting approach. Section 3.3 reports the estimated parameters of the observation model: MUA spatial profile *A*_*MUA*_ and CSD spatial profile *A*_*CSD*_. In Section 3.4, we finally examine the fitted parameters of the cortical column model and assess their similarity across the four recording sites (electrode penetrations).

### 3.1 Explanation of the target data

The waveforms for measured and predicted MUA and CSD for all 4 recording sites are shown in Fig. 6. These recording sites were selected to cover a wide range in tonotopic space in A1. More detailed depictions, alongside the goodness of fit (*R*^2^), are given in Fig. S4. The target data for model fitting includes the 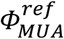 and 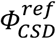 of the evoked response to the best frequency (BF) and non-BF tones (see Fig. 6A and B). The BFs of the four recording sites were determined to be 300 Hz, 1000 Hz, 500 Hz, and 6400 Hz, respectively. In 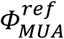, L4 (centering around Ch 10) shows higher MUA in BF conditions than in non-BF conditions. This corresponds to the frequency selectivity (tuning curves) of neural population responses that give rise to the tonotopic organization of A1. The BF response may include sustained activity (e.g., 500 Hz at site 3) or a second peak after 150 ms (e.g., 300 Hz at site 1). The non-BF responses at recording site 1 (i.e., 200, 260, 400, and 600 Hz) show negative firing rates (after subtraction of the baseline), reflecting the strong effect of lateral inhibition. In 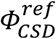, there are early L4 sinks and sources (especially clearly seen at sites 2 and 4) followed by sinks and sources at L2/3 and L5/6. The CSD also revealed longer-lasting sinks (red) and sources (blue) occurring in various channels after 100 ms (especially clearly seen at sites 3 and 4).

**Figure 6.**
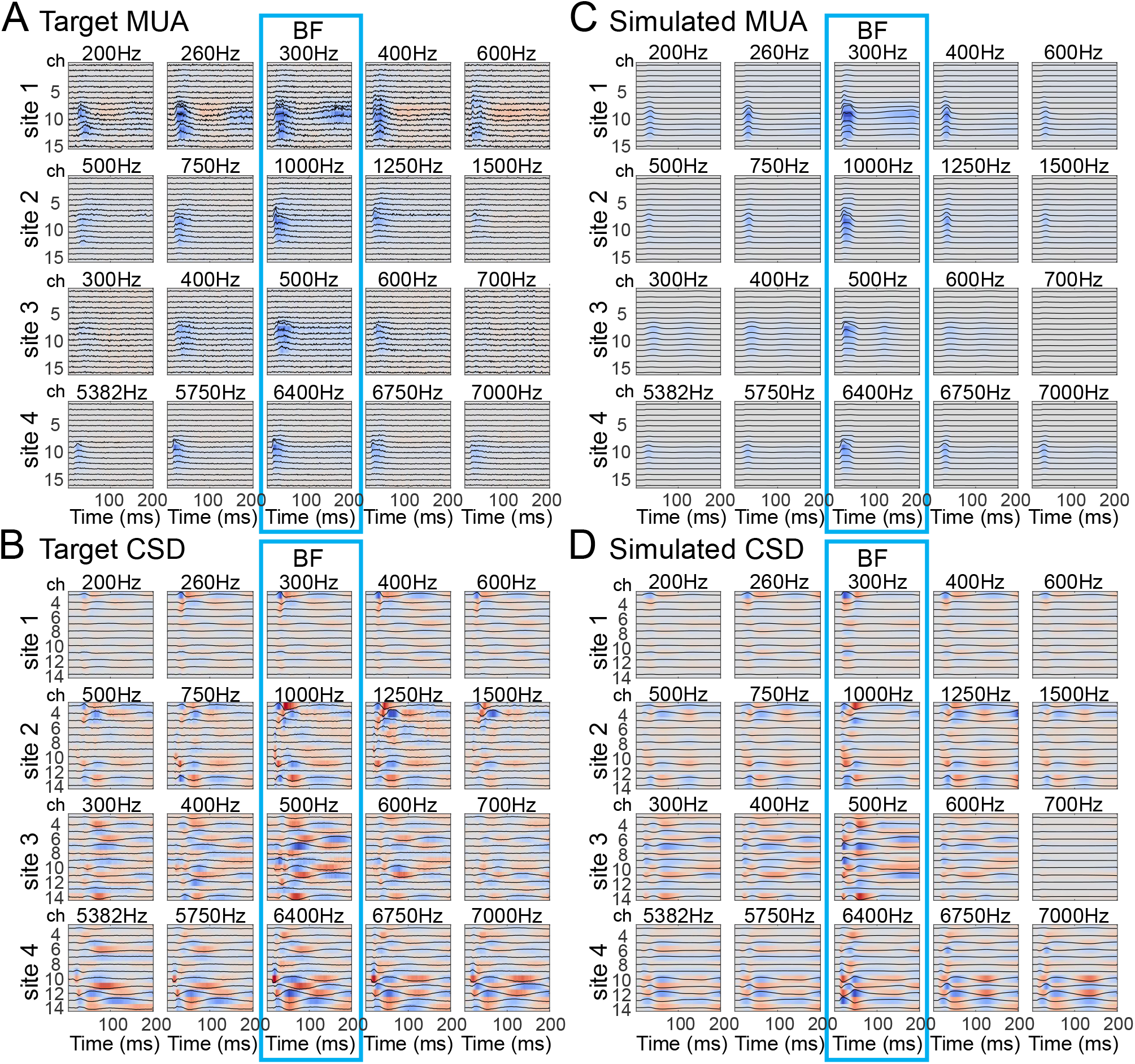
Visual comparison of target data and model simulations. (A, B) Target MUA (A) and CSD (B) at the four recording sites. The rows show the responses to different tone frequencies at each recording site. For example, the 1st row shows responses to 200, 260, 300, 400, and 600 Hz at site 1. The middle column (highlighted by the blue rectangle) represents the responses to tones at the BF of each recording site (300, 1000, 500, and 6400 Hz, respectively). The other columns represent non-BF responses. n the target data, the maximum value of each row was normalized to 1. (C, D) The simulated MUA (C) and CSD (D) from the final model solutions. Response amplitudes are color-coded (blue: positive; red: negative).

The predicted data *Φ*_MUA_ and *Φ*_CSD_ capture the general pattern of target data (Fig. 6C and D). The negative firing rate (relative to the pre-stimulus baseline, as at site 1) cannot be reproduced because we only allow non-negative firing rates and a non-negative MUA spatial profile to avoid overfitting (see Discussion). There are some other minor differences between the simulated and target responses. For example, the sustained firing rate (e.g., 500 Hz at site 3) is not well captured by the model. The patterns in *Φ*_*MUA*_ and *Φ*_*CSD*_ tend to be sharper and do not capture the smooth propagation across layers in 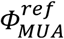, and 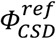, which is due to the limited number of populations (*N*_*pop*_ = 7) in the model.

### 3.2 Estimated latent variables

#### 3.2.1 Neural activity

The final model solutions provide estimated network dynamics that may underlie the empirical observations. Fig. 7 shows the firing rates *S*_*rate*_ of Column 1 (representing the recording site) as well as the strength of thalamic inputs based on the common solutions. The network dynamics across the recording sites, especially sites 1, 2, and 4, share many similarities. The thalamic inputs are stronger under the BF than the non-BF conditions, which agrees with the tonotopic organization of A1. The direct input connections lead to stronger early peaks in E and PV activity under the BF than the non-BF conditions (early intra-column E→PV effect). In contrast, the early peaks in SOM activity are often weaker under the BF than the non-BF conditions. This is due to the excitatory input from Column 2 via E→SOM connections (lateral inhibition, late inter-column E→SOM effect). Additionally, strong SOM activity at longer latencies appears to coincide with an increase in short-term facilitation of inter-column E-to-SOM connections (Fig. 7). The BF and non-BF responses show opposite dynamic patterns: strong PV inhibits SOM in BF responses, whereas the SOM inhibits PV in non-BF responses.

**Figure 7.**
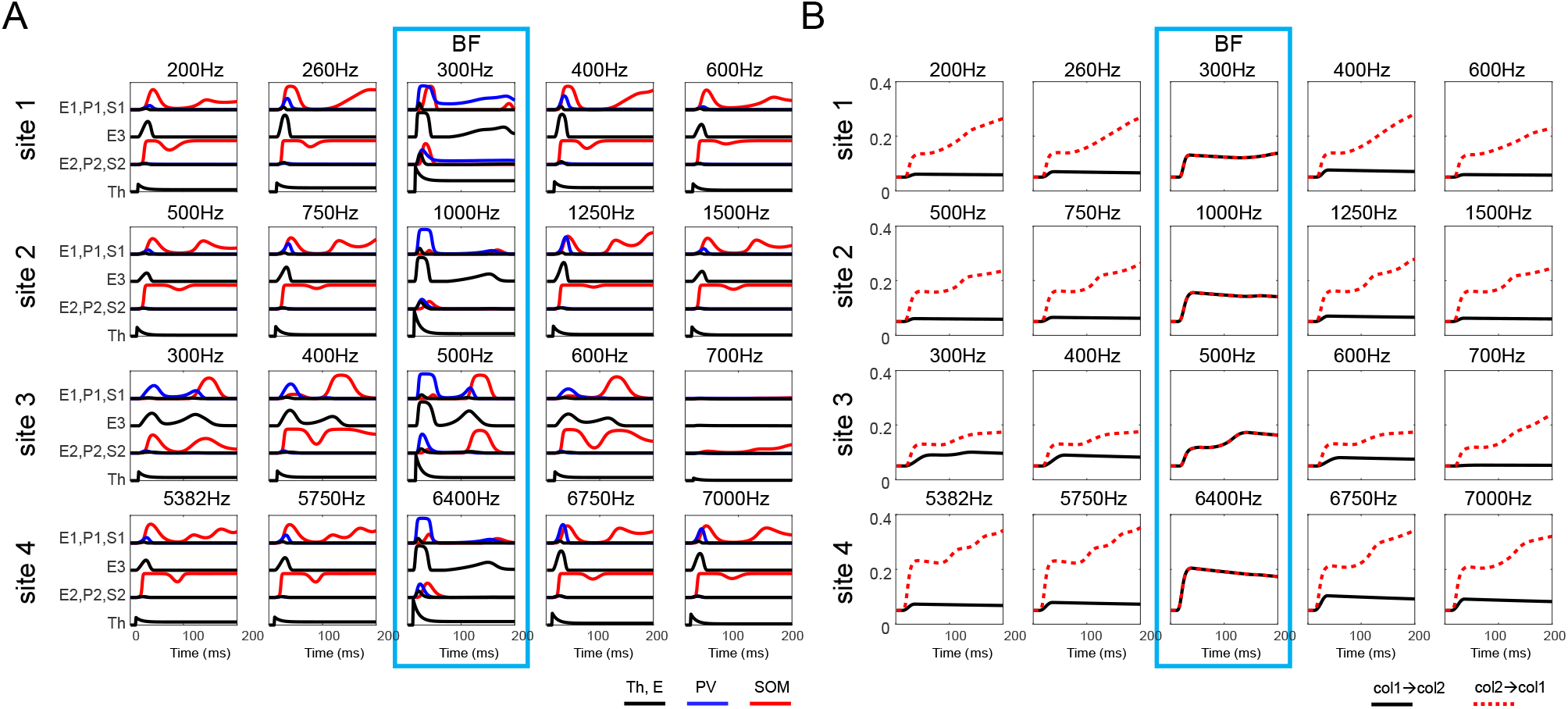
Estimated firing rates and plasticity of inter-column connections under BF and non-BF conditions. (A)The estimated firing rates are grouped into layers for visualization. The PV populations (blue curves) show stronger activity in BF responses (middle column) than in non-BF responses (other columns), where the peak values seem mainly to be affected by the strength of thalamic input (i.e., tonotopy). In contrast, the SOM populations (red curves) show long-lasting activity in non-BF responses, which reflects the effect of lateral inhibition. (B) The plastic lateral inhibition is reflected by the short-term plasticity variables x_inter−SE_ · u_inter−SE_ on the inter-column E-to-SOM connections W_inter−SE_ as in Eq. (9). The lateral inhibition from column 2 (BF site) to column 1 (recording site) is rising over time (red dashed curves) under non-BF conditions, whereas the lateral inhibition from column 1 to column 2 is slightly falling (black curves). In the BF conditions, the two curves are identical. (E1: L23 E population, P1: L234 PV population, S1: L234 SOM population, E3: L4 E population, E2: L56 E population, P2: L56 PV population, S2: L56SOM population, and Th: thalamic input)

#### 3.2.2 Contribution to EEG MEG components P1, 1, and P2

The equivalent current dipole (ECD) *Φ*_*ECD*_ derived at each recording site provides a link between intracranially-recorded LFPs and extracranially-recorded EEG/MEG signals. The ECD can be calculated by Eq. (14), where the current flows associated with activity of specific neural populations contribute to the ECD by different weights and directions based on the CSD spatial profile *A*_*CSD*_. Fig. 8A shows the predicted ECDs and the contributions of current flows by the E, PV, and SOM populations at the four recording sites under BF and non-BF conditions. Although there is no ground truth for verification, the predicted ECDs (the black curves in Fig. 8A) show a pattern of positive/negative/positive deflections at approximately 25, 40, and 80 milliseconds (site 1 deviates from that with considerably longer latencies) that can be related to the EEG/MEG components P1, N1, and P2. These deflections are well-known to occur in humans in response to pure tone stimulation at latencies of approximately 55, 100, and 160 milliseconds, while corresponding peaks have been described in macaques at 30, 55, and 70 milliseconds (Itoh et al., 2019). The deflections are the net result of the summation and cancellation of current flows. To gain insight into cell-type specific contributions to these deflections, we summed the dipole magnitudes from each of the cell types (while keeping the thalamic input separate), resulting in only four dipole magnitudes (i.e., 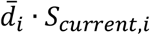 where *i* ∈ {*E, PV, SOM, Th*} as in Eq. (14)). For example, the PV dipole signals (yellow curves in Fig. 8A) represent the gross current flow in pyramidal dendrites (i.e., the IPSP on E1, E2, and E3 populations) elicited by the PV neurons (i.e., PV1 and PV2 populations).

**Figure 8.**
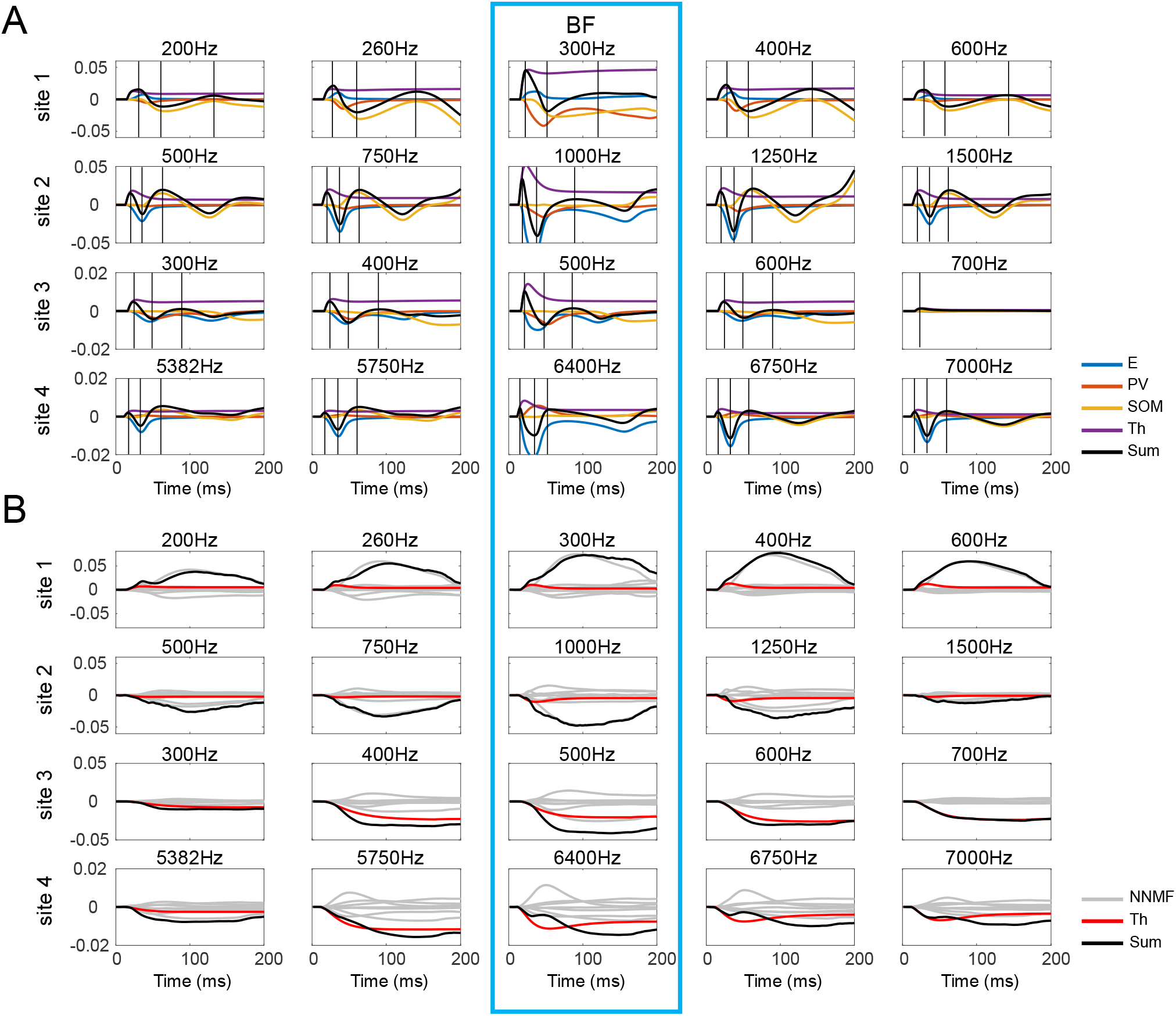
Simulated equivalent current dipole (ECD) signals. (A) The simulated ECD (black curves) reflects the contribution of a recording site to the EEG/MEG signal (before considering the orientation with respect to the MEG sensors). The simulated ECD signal is calculated as the sum of population-level CD signals (colored curves) representing the effective current flows along the long dendrites of pyramidal cells induced by individual neural populations or thalamic input (E: E1, E2, and E3; PV: PV1 and PV2; SOM: SOM1 and SOM2; Th: thalamic input). The vertical lines indicate the deflections of the ECD signals. (B) Simulated ECD signals obtained by the NNMF approach.

In Fig. 8A, we find that the first positive peak of the ECD (related to P1) is consistently due to the thalamic input (purple curves). The first negative peak of the ECD (related to N1) can be due to either the E current (blue curves, sites 2, 3, and 4) or the PV current (red curves, site 1), which depends on the dipole direction derived from the CSD spatial profile. The second positive peak of the ECD (black line), associated with the P2, is not uniquely attributable to a particular deflection in the cell currents. For the non-BF condition, it mostly coincides with a positive deflection of the SOM current, except for site 3, where it coincides with a peak in the E current. For the BF condition, the P2 peak is mostly associated with a peak in the E current, except for site 1, where we see a positive peak in the PV current at that latency. Based on these predicted ECDs and cell-type specific contributions, the P1 component is likely to result from the thalamic input (including the BF and non-BF columns). The N1 component is likely to result from the activity of E/PV neurons (also including the BF and non-BF columns). The origin of the P2 component cannot be identified so clearly. It may be caused by a change in SOM activity for non-BF columns (not confirmed in site 3) and by a change in E activity in BF columns (not confirmed in site 1).

The model fitting approach provides a way to decompose the LFP into temporal and spatial components and predict the ECD at the recording site. We were curious whether a blind decomposition approach would yield similar predictions. To examine this possibility, we decomposed the MUA using non-negative matrix factorization (NNMF), determined optimized current flows and CSD spatial profile, and calculated the dipole signal (Section 2.5). We found that the NNMF approach, although it leads to improved goodness of fit (Fig. S5) compared with the model fitting approach, yields implausible time courses of the ECDs (Fig. 8B). The time courses of the magnitude of dipoles underlying auditory evoked responses usually feature a pattern of alternating peaks and troughs following that of the respective scalp recordings (see, e.g., Knösche et al., 2002). As the modeling approach does reproduce an EEG/MEG time course (through the ECD) with the typical peaks of P1, N1, and P2, it appears to be a promising candidate for realistically accounting for the MUA and CSD. Note that the precise latencies of these peaks are usually quite variable even in humans and depend on a number of factors (Picton, 2010).

#### 3.2.3 Parameters of the observation model

The observation model contains an MUA spatial profile *A*_*MUA*_ and a CSD spatial profile *A*_*CSD*_ that project firing rates *S*_*rate*_ and current flows *S*_*current*_ to observations *Φ*_*MUA*_ and *Φ*_*CSD*_. We now check the estimated spatial profiles A_MUA_ and A_*CSD*_ provided by the final solutions. The MUA spatial profiles *A*_*MUA*_ represent the sensitivity of the 16 laminar electrode channels to the firing rates of neural populations. Fig. 9A shows the MUA spatial profile at each recording site (electrode penetration). The distributions of E3 center around channel 10, which is consistent with the distribution of L4 excitatory neurons. However, the distributions of E1 and E2 form one to two clusters, a scenario which does not agree with the expected distribution of excitatory neurons in L2/3 and L5/6. This suggests that the definition of neural populations, after model fitting, do not strictly adhere to the initial laminar definition (i.e., the default connectivity) but changes based on the temporal dynamics of neural activity. The distributions of E3 center around channel 10, which is consistent with the distribution of L4 excitatory neurons. However, the distributions of E1 and E2 form one to two clusters, which do not match the distributions of excitatory neurons in L2/3 and L5/6, as they were assumed in the definition of the model. Similar observations are made for SOM and PV neurons. This demonstrates that the best grouping of neurons into a few masses, such that their mean activities explain the measured effects, does not necessarily coincide with their strict assignment to some layers.

**Figure 9.**
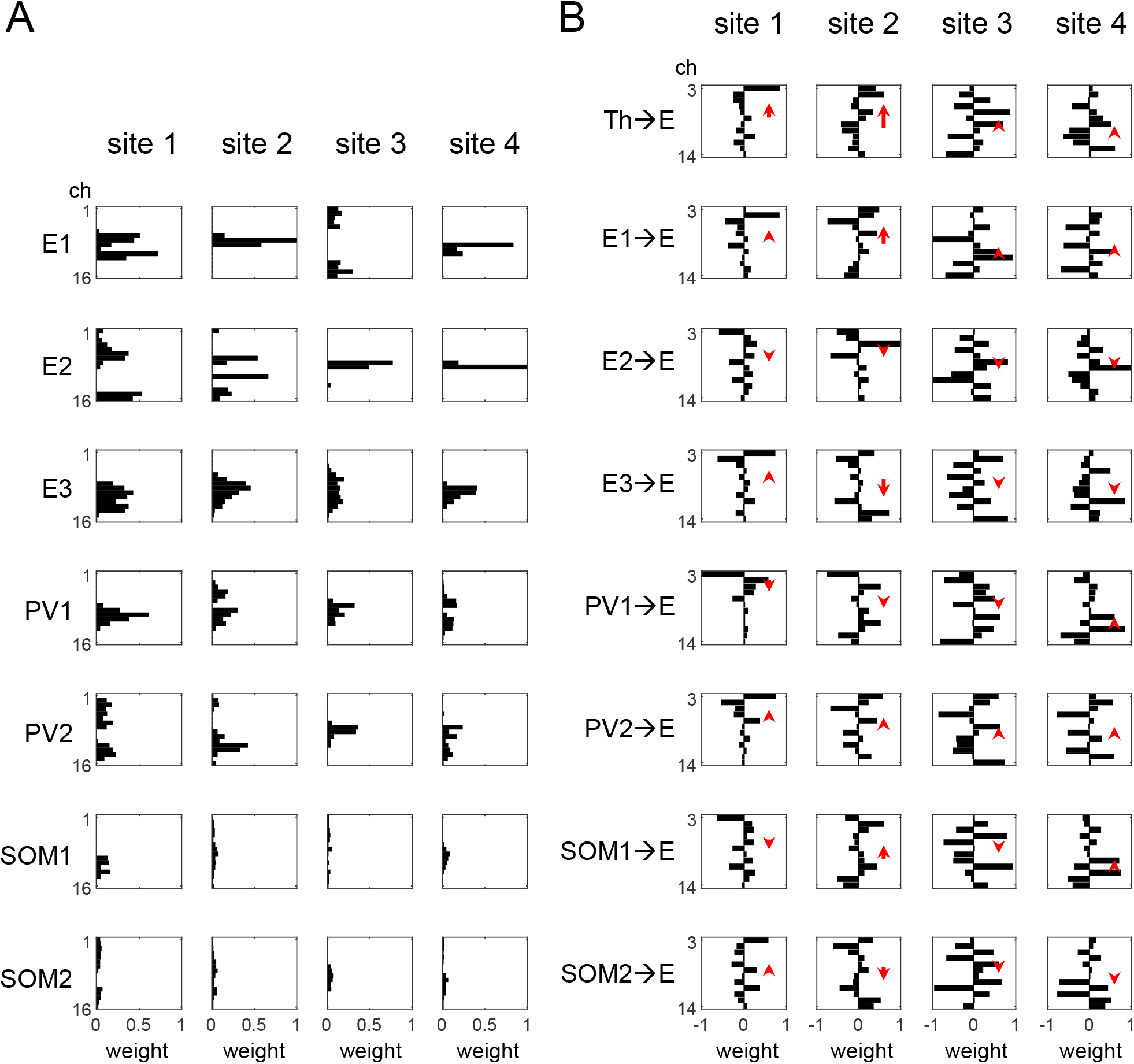
Estimated spatial profiles. (A) MUA spatial profiles. Each column represents the laminar distribution (across electrode channels) of firing rates of neural populations within each recording site. (B) CSD spatial profiles. Each column represents the corresponding laminar distribution of sinks and sources. The red arrow indicates the dipole length and direction (from center of sinks to center of sources).

Fig. 9B shows the CSD spatial profile of each recording site, which represents the overall distribution of current sinks and sources along the dendrites of the excitatory cells (incl. E1, E2, and E3). We found that the distributions of sinks and sources interlace and spread widely, and differ across recording sites. This may correspond to the relatively high variability in CSD across recording sites. This may be also due to the fact that the depth of the 16-channel electrode within the cortex is not necessarily identical across the 4 electrode penetrations.

### 3.3 Fitted parameters

Next, we checked whether the common solutions for the four recording sites share similar patterns. In Fig. 10, we show parameters related to the tuning curve and lateral inhibition. The tuning curve is characterized by four parameters (*θ*_25−28_) which define the strength of thalamic input to the recording site (Column 1) under the four non-BF conditions. The tone frequencies in Fig. 10 are plotted on a log scale. The lateral inhibition is characterized by five parameters (θ_20−24_) which rescale the default strengths of inter-column E-to-SOM connections. We found M-shape lateral inhibition curves (Fig. 10, lower panel), where lateral inhibition increases as the tone frequency deviates from the BF tone but decreases as the tone frequency deviates further (not reached for site 2). The input strengths under the four non-BF conditions are all weaker than 0.5, and decrease as the tone frequency becomes more distant from the BF of the neurons (Fig. 10, upper panel), which is consistent with features of spectral tuning curves in A1. The lateral inhibition and tuning curves directly affect the activities of PV and SOM populations, respectively, as demonstrated by the firing rates under BF and non-BF conditions in Fig. 7.

**Figure 10.**
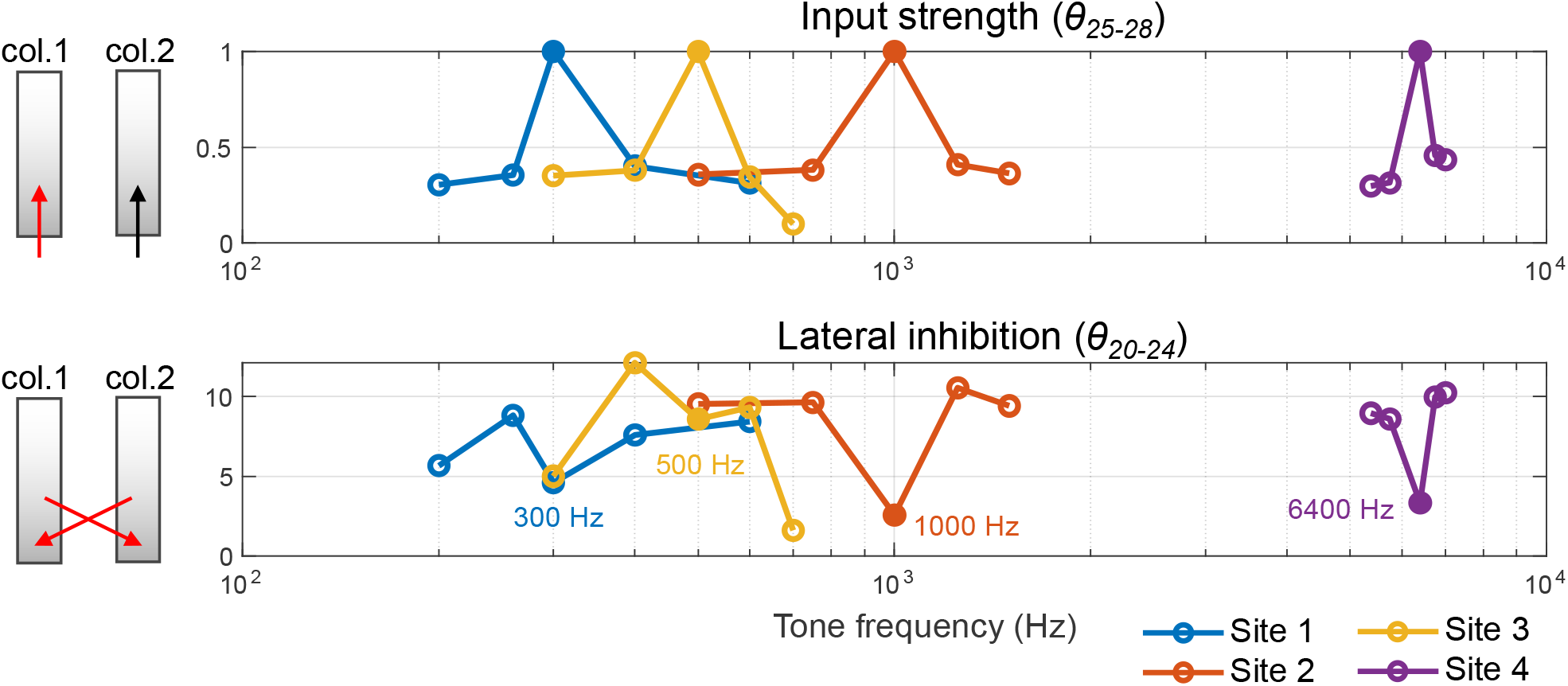
nput strength and lateral inhibition in evoked responses. The fitted parameters are plotted against tone frequency (Hz in log scale). The parameters θ_25−28_ rescale the default thalamic input strengths I_1−4_ (as illustrated by the red arrow to the cortical column 1, upper left panel) under the four non-BF conditions (depicted as open circles). Note that the input strength to Column 1 under the BF condition (solid circles) is always fixed to 1. The parameters θ_20−24_ rescale the default lateral inhibition W_inter−SE_ (as illustrated by the red arrows between the two cortical columns, lower left panel) under the BF conditions (solid circles) and four non-BF conditions (open circles). The BFs corresponding to the four recording sites (300, 1000, 500, and 6400 Hz) are indicated alongside the solid circles.

We then checked the cost function surface by separately scanning each parameter around the solutions (scanning ranges as listed in Table 6). The surfaces of the four recording sites are shown in Fig. 11. In general, we found that the values of solutions (indicated by triangles) are close to each other in the scan range, and the cost (normalized SSE) increases as the values deviate from the solutions. Hence, at least local minima of the cost function exist, indicating clearly defined solutions. The fact that these solutions are close to each other in parameter space adds to their credibility, based on the assumption that the wiring schemes of cortical columns are similar within the auditory cortex.

**Figure 11.**
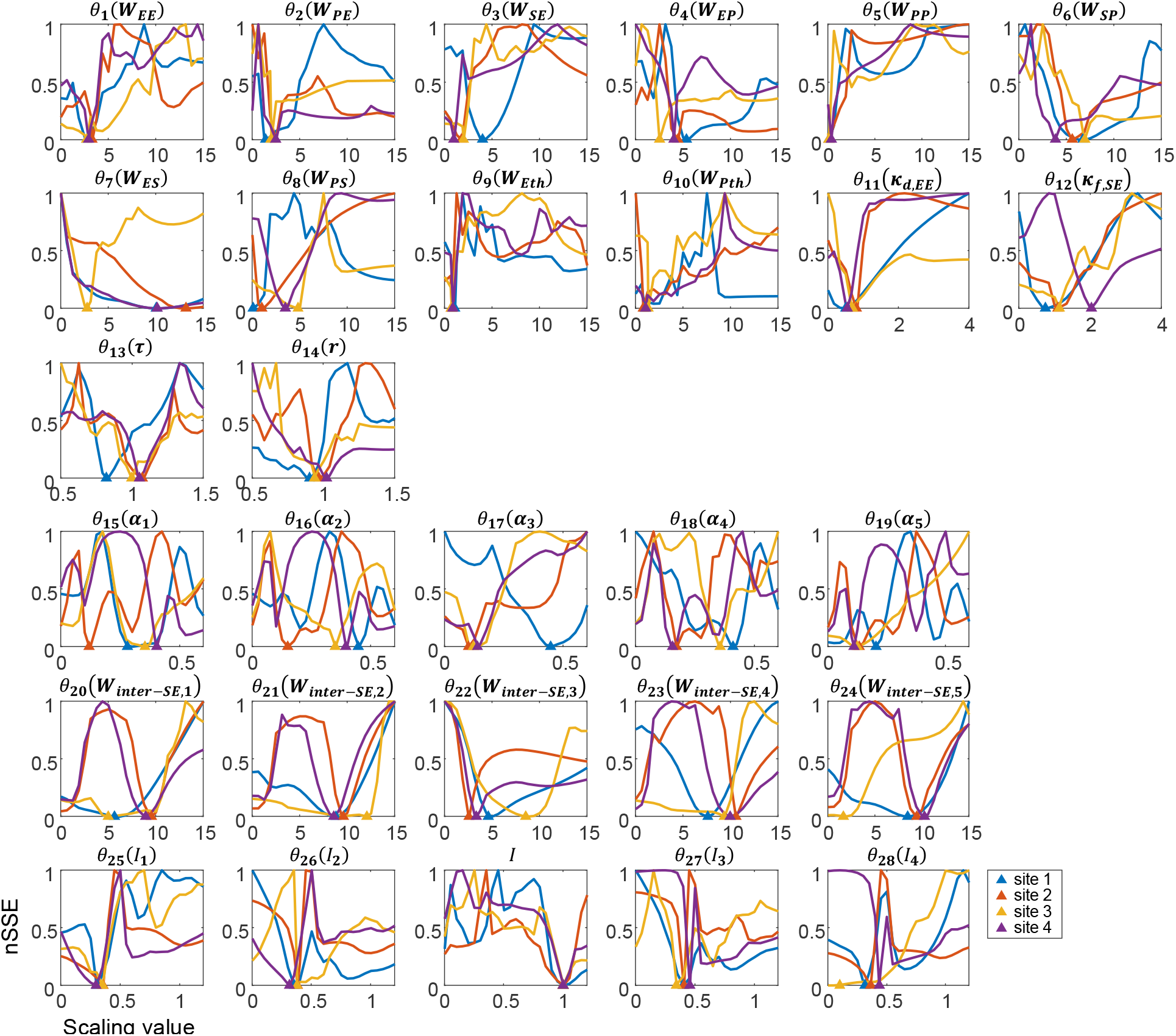
Normalized sum of squared errors (nSSE) between predicted and target observations as function of the free parameters. Each plot shows how the nSSE varies as a function of one of the free scaling parameters, while the rest of the parameters are kept constant to their optimal values. These scaling parameters (column 3 of Table 6) are stated on top of each plot together with the scaled model parameter (column 1 of Table 6) in brackets. See Table 6 for an overview of the model parameters, their default values, the free parameters scaling them, and the scaling ranges. The x-axes represent the parameter values normalized to their respective default values. The triangles represent the solutions for the four recording sites.

## 4 Discussion

In this study, we combined a detailed neural circuit model of the cortex with fine-grained laminar LFP recordings in monkey primary auditory cortex (A1) to estimate cell-type specific contributions to both intracranially- and extracranially-recorded signals. We show that evoked responses at four example recording sites, covering a wide range in tonotopic space in A1, share similar network dynamics (i.e., E, PV, and SOM activity) (Fig. 7), but can show diverse patterns in net transmembrane extracellular current flow, as reflected by CSD analysis, due to the variation in spatial profiles (Fig. 9). The four recording sites also share similar input curves and lateral inhibition curves (Fig. 10) as well as similar intra-column configurations (Fig. 11). These results support the notion of canonical microcircuits (Douglas et al., 1989; Douglas & Martin, 1991, 2004a) and a consistent pattern of neural dynamics and interactions contributing to sensory processing in A1.

We demonstrate the feasibility of our model-fitting approach by transforming laminar profiles of MUA and CSD into products of spatial and temporal components. The fitted model provides insights into neural interactions and cell-type specific activities contributing to equivalent current dipoles (ECDs) underlying typical EEG or MEG recordings (Fig. 8A). This is supported by the plausible ECD signal derived from the estimated current flows and CSD spatial profile. In contrast, an alternative approach based on non-negative matrix factorization, despite a higher goodness of fit to MUA and CSD response profiles (Fig. S5A), is relatively uninformative with regard to elucidating the activities of distinct neural populations (Fig. S5B) and plausible synthesized ECD signals (Fig. 8B).

### 4.1 PV-SOM interaction

We observed distinct patterns in the estimated activity of PV and SOM interneurons between BF and non-BF responses (Fig. 7). In BF responses, PV interneurons show faster and stronger activity than SOM interneurons. In non-BF responses, PV interneurons show relatively weak activity, and the activity of SOM interneurons dominates after around 50 ms. This phenomenon results from various literature-based settings for PV and SOM neurons in the cortical column model with regard to thalamic input (no direct input to SOM, Ji et al., 2016), synaptic time constants (slow dynamic in SOM, Dura-Bernal et al., 2022; Jouhanneau et al., 2018), inter-column connections (only E-to-SOM, considering a wider spatial distribution of inhibitory innervation by SOM than PV neurons, Kato et al., 2017; Lakunina et al., 2020), and short-term plasticity (STF on E-to-SOM connections, Hayut et al., 2011; Silberberg & Markram, 2007). In other words, the early PV activity is directly related to tonotopic (thalamic) input, and the late SOM activity is directly related to lateral inhibition. Based on this distinction, the switch between the “ pattern” and “non-pattern” along the tonotopic axis is sharpened by mutual inhibition between PV and SOM interneurons (Hahn et al., 2022) and should be observable in possible future experiments involving detailed recordings from different neuron types, for example, through calcium imaging.

### 4.2 Generation of P1, 1, and P2 components

Auditory evoked responses recorded via EEG/MEG are typically characterized by a temporal sequence of positive and negative deflections or waves designated as the P1, N1, and P2 components. Studies based on the Human Neocortical Neurosolver (Kohl et al., 2022; Neymotin et al., 2020) suggest that the P1 and P2 components are partly generated by upward currents within the dendrites of cortical pyramidal neurons due to bottom-up inputs, while the N1 component is partly generated by downward currents associated with top-down inputs. Studies using a multi-column model of the auditory cortex (Hajizadeh et al., 2019, 2021, 2022) further emphasize the dependence of current orientation on the cellular location of the active synapses (e.g., apical/somatic), synaptic type (i.e., excitatory/inhibitory), inter-column connection type (e.g., feedforward, feedback, within-field), and folding of the cortex (i.e., the neuroanatomical topography of the cortical surface). At the level of a cortical column, our analyses of the cell-specific contributions to ECDs (Fig. 8A) suggests that initial thalamic input primarily contributes to P1, subsequent early activity of E and PV neurons (both BF and non-BF columns) primarily contributes to N1, and late SOM activity (especially in non-BF columns) joins the contribution to P2. We observed variability in peak latencies of the ECD signals at the four recording sites (Fig. 8A), due to the variance in thalamic inputs, intra-column connection strengths, and the CSD laminar spatial profile across cortical columns.

### 4.3 Patterns of activity propagation

In this study, we were unable to identify a consistent pattern of neural activity propagation across cortical layers and neural populations contributing to the auditory evoked response. Such patterns (e.g., from L4 to L2/3 and to L5) have been assumed in canonical cortical column models of visual and somatosensory areas (e.g., (Douglas & Martin, 2004b)). However, such a stereotypical pattern might not always be the rule. For example, a study using thalamic stimulation suggested that activity in supragranular layers is initiated by infragranular cells and regulated by feed-forward inhibitory cells (Krause et al., 2014). Moreover, specific patterns of information propagation are less likely to be found from a more detailed and complex network derived from existing animal-model databases (Billeh et al., 2020; Campagnola et al., 2022; Ji et al., 2016; Markram et al., 2015). Based on our estimated firing rates (Fig. 7) and MUA spatial profile (Fig. 9A), the three E populations (E1, E2, and E3) show early peaks at similar latencies because they all receive thalamic input. Clear propagation from L4 to L2/3 to L5 is therefore not observed. As for the activity of inhibitory neurons, PV1 activity is in general stronger than PV2 activity in BF responses, and SOM2 activity is stronger than SOM1 activity in non-BF responses. However, this pattern cannot be projected onto specific cortical layers, because the estimated MUA spatial profiles of the same cell type (e.g., PV1 vs. PV2; SOM1 vs. SOM2) are not spatially exclusive. While our cortical column model was built with default connectivity based on laminar classification, the neural populations are found to spatially overlap after model fitting. Such spatial overlap of functional components may provide an alternative framework for understanding the neural computations underlying sensory processing, for example, the role of lateral inhibition in A1 in rhythmic masking release (Fishman et al., 2012), spectral resolution (Fishman & Steinschneider, 2006), and complex tone processing (Fishman et al., 2000b).

### 4.4 Limitations and future directions

Several important limitations of the present work should be noted. First, our model considers only thalamic and lateral input and disregards inputs from areas outside of primary auditory cortex, e.g., frontal and entorhinal cortices (Schaefer et al., 2015). Such input may be increasingly important for explaining neural activity occurring at longer latencies. Second, the firing rate in our model ranges from 0 to 1, corresponding to baseline activity and maximum firing rates of each cell type, respectively. We restricted the sigmoid functions to be non-negative in order to prevent unreasonable cancellation between negative and positive firing rates in the fitting procedure. This comes with the price of failing to explain negative (relative to the pre-stimulus baseline) MUA activity. Third, in order to avoid over-fitting, we reduced model complexity by making a number of simplifying assumptions. For example, the intra-column settings of the two columns are assumed to be identical. The ratio of thalamic input to E and PV neurons is assumed to be fixed. The inter-column connections are assumed to be symmetric. The MUA spatial profile is assumed to have a fixed ratio of sensitivity to E, PV, and SOM neurons. In general, these assumptions lead to a reduced goodness of fit. We cannot guarantee that we have found globally optimized solutions, but current best solutions show consistent neural activities and fitted parameters across the four recording sites examined. The current best solutions can serve as a prior to improve goodness of fit to data from individual sites using a more flexible model, or one with a greater number of free parameters.

In this study, we demonstrated the feasibility of our model-fitting approach in estimating cell-type specific activity across cortical layers based on LFP recordings in A1. So far the target observations only include tone-evoked responses (BF and non-BF conditions) at four recording sites. Moreover, our cortical column model was designed with a relatively simple architecture in order to avoid an underdetermination issue arising from insufficient data constraints. However, our simplified model could be extended in several ways. For example, VIP interneurons could be included to examine the modulatory effects of attention on different inhibitory states (Hahn et al., 2022). The matrix thalamic input (innervating the supragranular cortex) could be included as tone-insensitive input (Müller et al., 2020) to investigate neural mechanisms underlying other neurophysiological phenomena in auditory cortex, such as “Off” responses and mismatch responses (e.g., (Fishman, 2014; Fishman & Steinschneider, 2009)). Corticothalamic pathways (which emanate from L5/6 to the thalamus) could also be included to model the modulation of thalamic input (Antunes & Malmierca, 2021). Lastly, spontaneous activity in LFP recordings could be considered in future work to examine the neural underpinnings of interactions between spontaneous and stimulus-evoked neural activity in auditory cortex (e.g., Dura-Bernal et al., 2022).

## 5 Conclusions

We demonstrated the feasibility of a model-fitting approach in estimating cell-type specific population activity from laminar LFPs evoked by sound stimuli, as well as estimating cell-type specific contributions to the P1, N1, and P2 components of extracranial evoked responses. This model-fitting approach provides a foundation for further investigation into the neural dynamics of cortical sensory processing.

## Code availability statement

The main codes used in this article can be found at https://github.com/vscChien/LFPmodeling.

## Acknowledgments

Neurophysiological data were obtained in collaboration with Dr. Mitchell Steinschneider (funded under NIH Grant DC00657), with invaluable technical support provided by Jeannie Hutagalung and Shirley Seto.

## Supplementary figures

**Figure S1.**
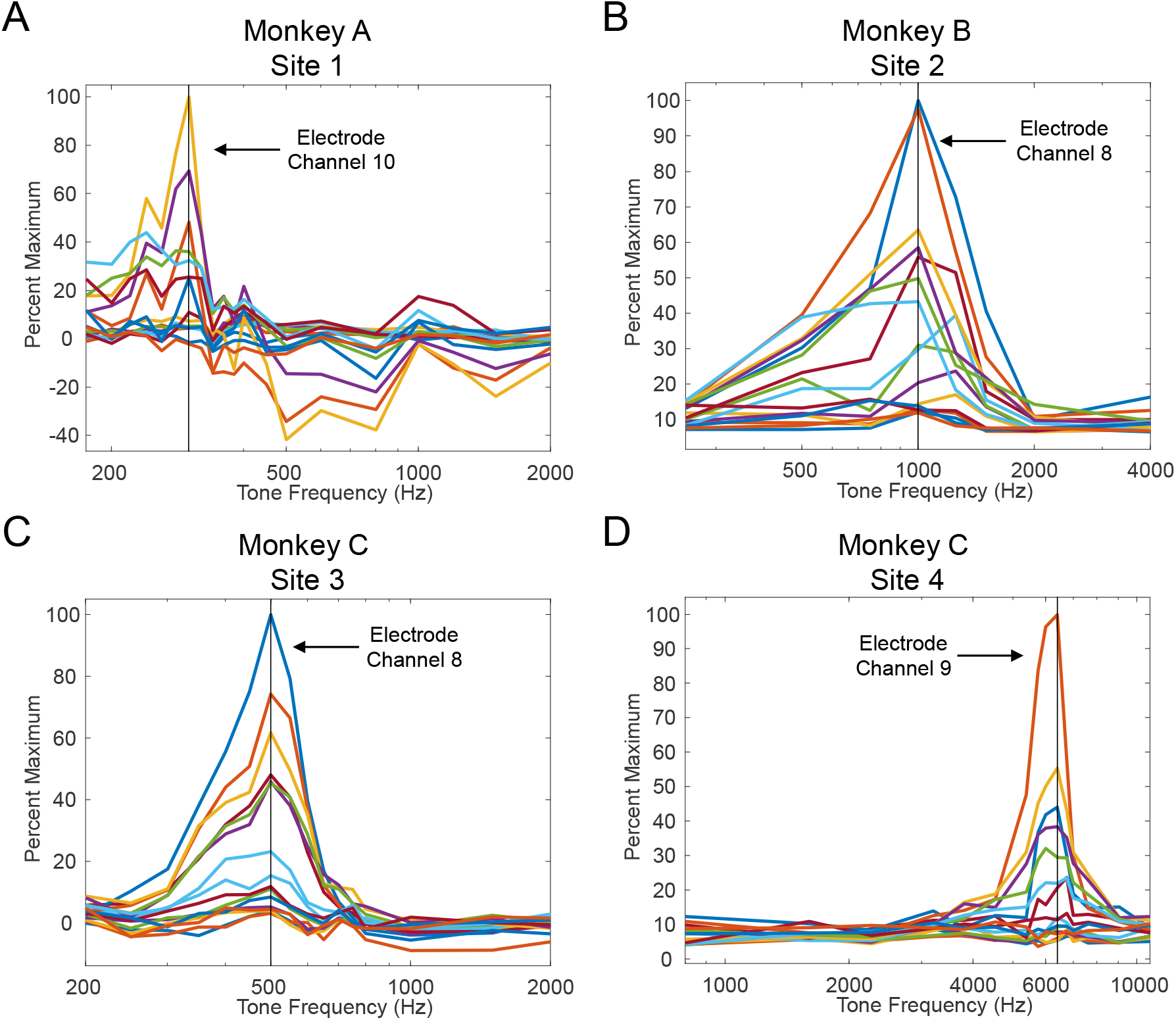
Neural population tuning curves obtained at each of the 16 electrode channels at the 4 recording sites (electrode penetrations) analyzed in this report (represented in panels A-D, respectively). Each colored plot represents the tuning curve of MUA at each electrode channel spanning the laminar extent of A1 (plots have been normalized to the maximum-amplitude response). The tuning curve for the electrode channel at which the maximal-amplitude MUA was recorded is indicated by the arrow. Note that this channel is generally located in the middle of the electrode array and positioned within the middle laminae of A1. The Best Frequency (BF) corresponding to the peak of the tuning curve is indicated by the dashed vertical line. Note the high consistency in BFs across channels displaying the larger-amplitude responses, demonstrating that BF is relatively uniform across layers within a ‘cortical column’ in A1.

**Figure S2.**
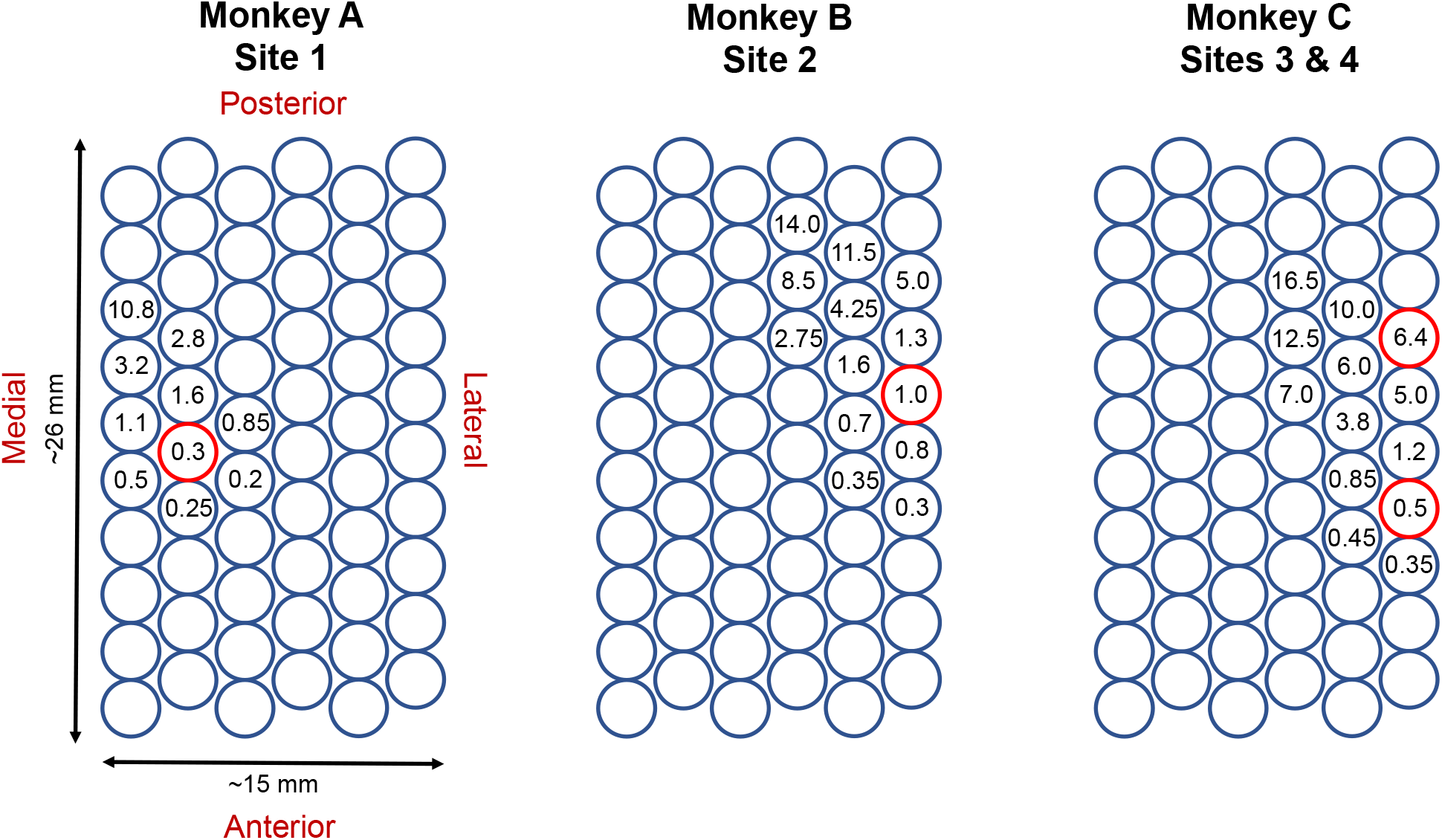
Schematic of matrices used to guide linear-array multi-contact electrodes perpendicularly into auditory cortex within the left hemisphere in Monkey A, B, and C, as indicated. Circles represent the tops of guide tubes into which the electrode was inserted during a given recording session. Numbers within the circles indicate the Best Frequency (BF; in kHz) of the neural population recorded in each electrode penetration, where a clear and unique BF could be determined based on pure tone responses. Red circles represent the 4 recording sites selected for analysis in the present report. These 4 sites were selected because they display stereotypical features of laminar AEP, MUA, and CSD neurophysiological response profiles which are characteristic of primary auditory cortex (A1). Note the approximate anterior-lateral to posterior-medial anatomical spatial gradient of low-to-high BF in all 3 animals, consistent with the tonotopic organization of A1 in macaques.

**Figure S3.**
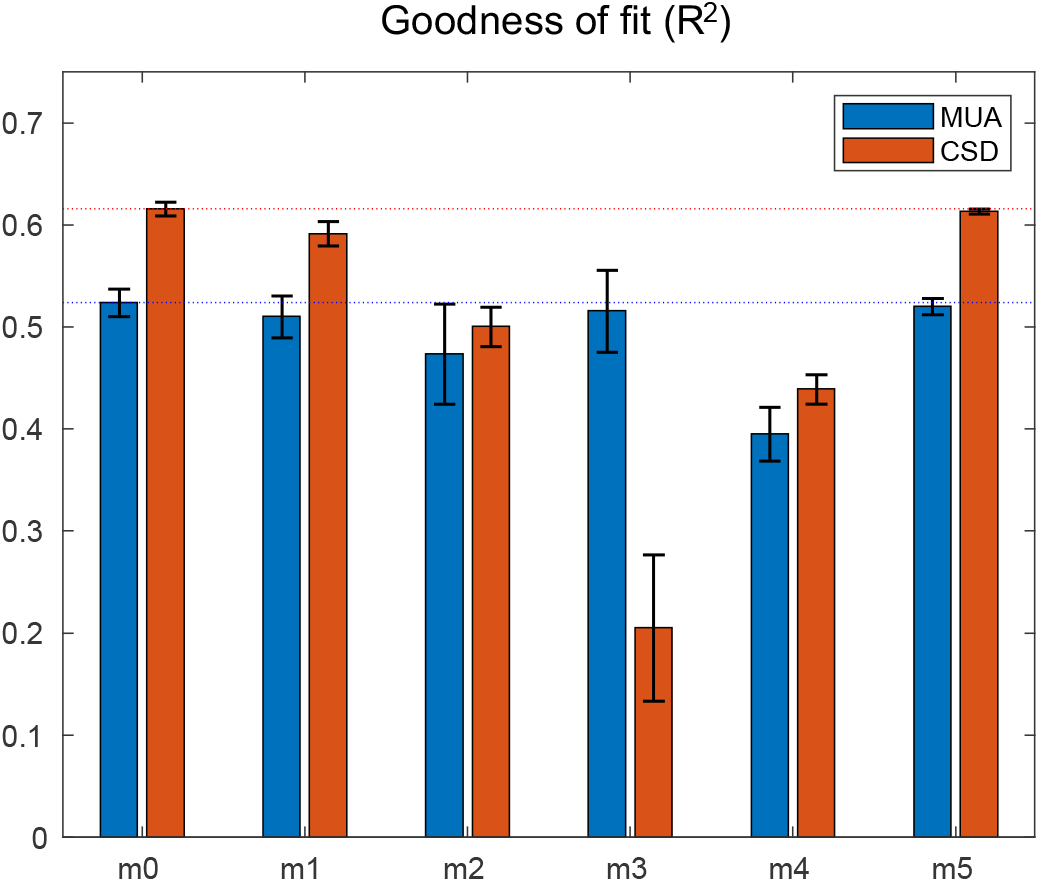
Performance of original model (m0), the reduced models (m1-4), and the expanded model (m5). To reveal the relative impact of thalamic input and inter-column connection on the fitting performance, we examine the goodness of fit (R^2^) by several variant models (four reduced models and one expanded model) on an exemplary recording site (site 2). The reduced models include m1: thalamic inputs only to L4, m2: no inter-column connectivity, m3: no PV populations, and m4: no SOM populations. Optimization procedures were performed six times for a stability check (standard deviation indicated by error bars). The results show that the reduced models m2-4 perform worse than the original model (m0), and the reduced model m1 has a similar performance. In the expanded model (m5), we include inter-column E-E connections between the L56 E populations, between the L23 E populations, and from the L23 E to L4 E population. The extra current flows by the inter-column E-E connections are included in producing the simulated CSD. Optimization procedures were performed six times with different initial values of inter-column E-E connections. The results show that m5 performs similarly to m0. The estimated firing rates of m5 (not shown here) are very similar to m0, suggesting that the original model (m0) suffices to characterize the network parameters and dynamics.

**Figure S4.**
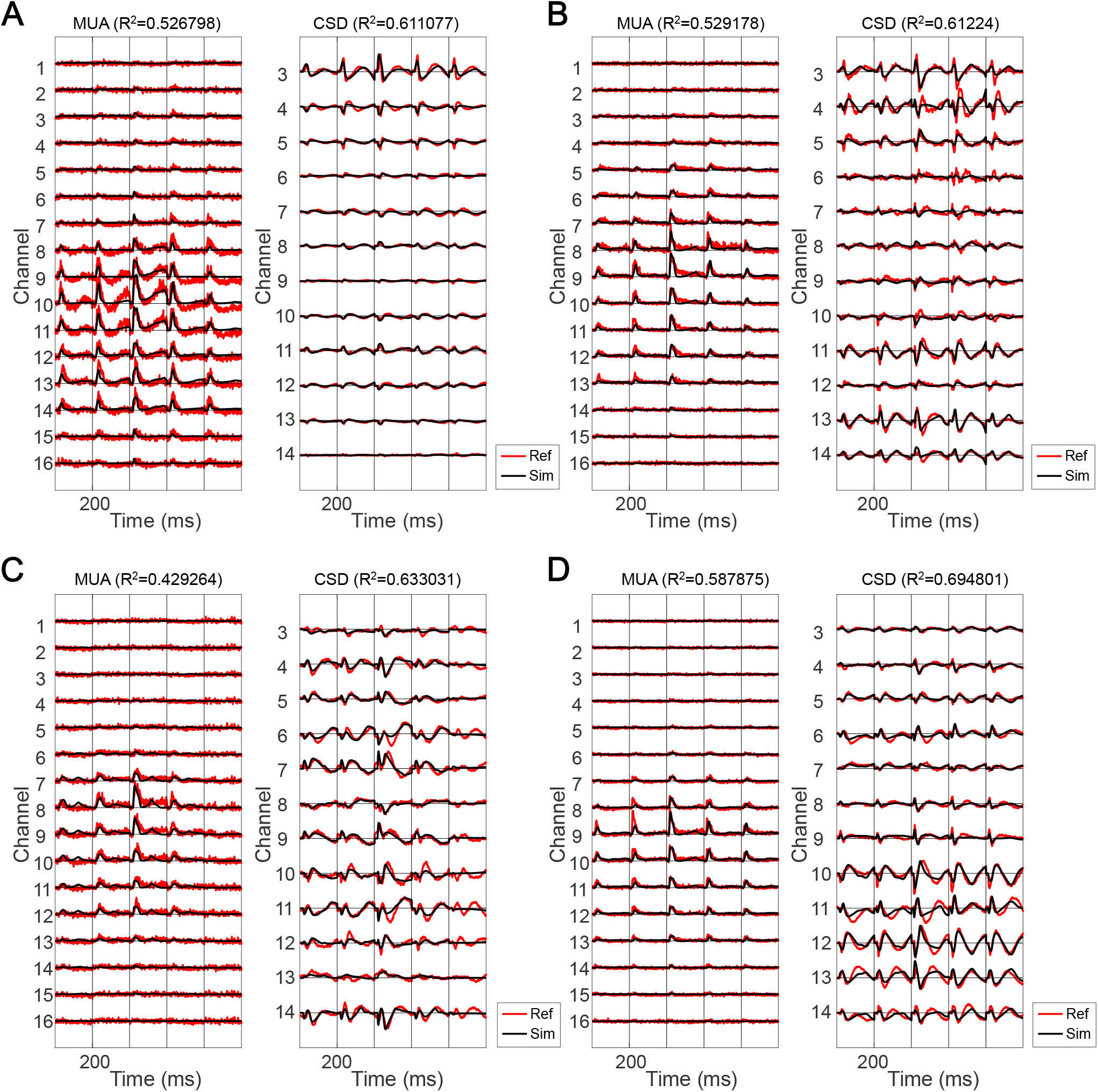
The target MUA and CSD (red) and the model simulations (black) at sites 1 to 4 (A-D).

**Figure S5.**
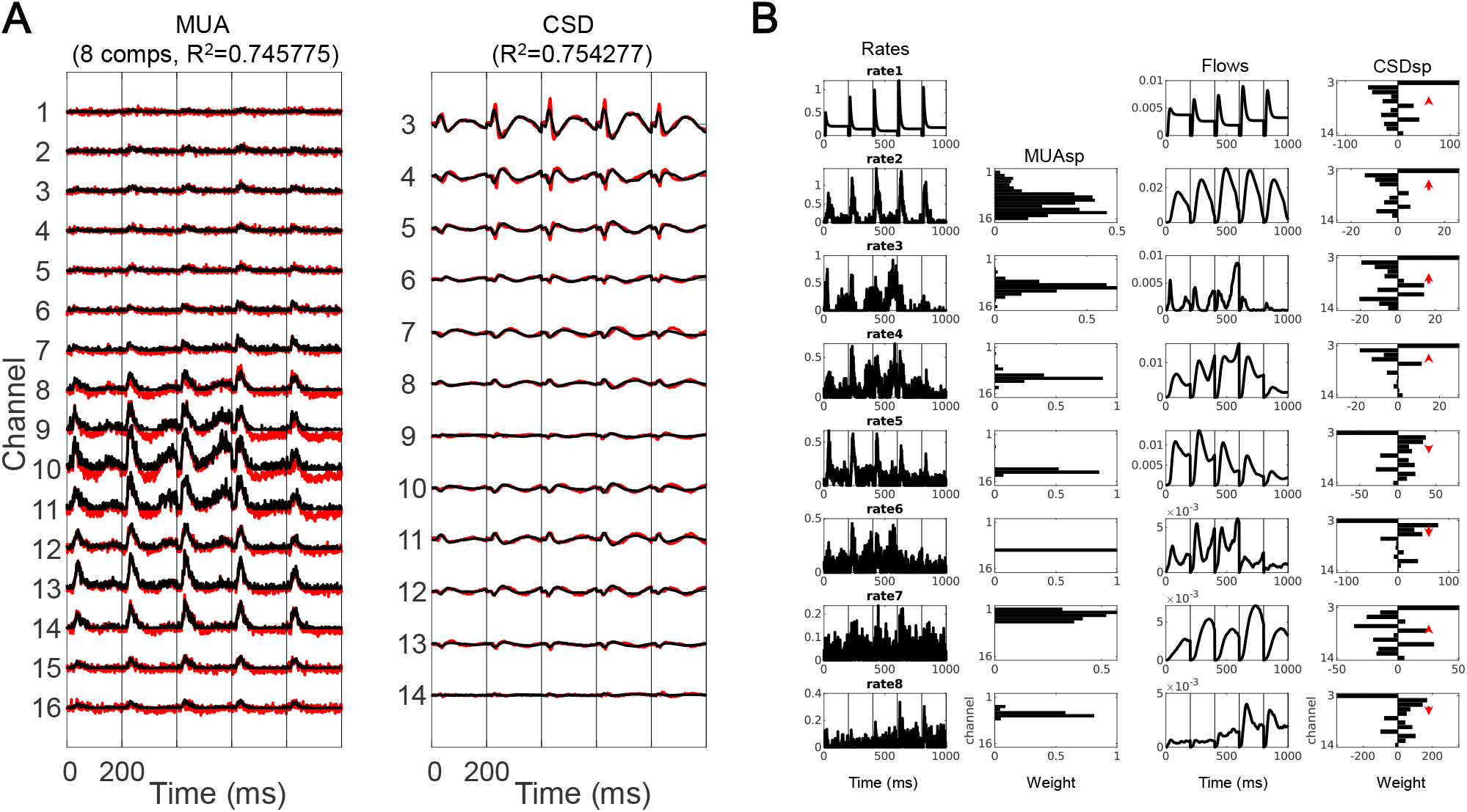
The NNMF decomposition approach (site 1 used as example). (A) The target MUA and CSD (red) and the model simulations (black). (B) The MUA is decomposed by NNMF into firing rates (rates 2 to 8 in 1st column) and MUA spatial profile (2nd column). The thalamic input (rate1 in 1st column) and the firing rates are convolved with alpha kernels to generate current flows (3rd column). The CSD spatial profile is the regression coefficients, and the centers of sinks and sources are shown in red arrows (4th column).

**Table S1A.** Connection probabilities after averaging across layers.

| To \ From | E1 | E2 | E3 | PV1 | PV2 | SOM1 | SOM2 |
| --- | --- | --- | --- | --- | --- | --- | --- |
| E1 (L2/3) | 0.1600 | 0.0105 | 0.1400 | 0.3305 | 0.0500 | 0.3370 | 0.0845 |
| E2 (L5/6) | 0.0415 | 0.0503 | 0.0680 | 0.0395 | 0.1713 | 0.0118 | 0.1192 |
| E3 (L4) | 0.0160 | 0.0035 | 0.2430 | 0.2435 | 0.0500 | 0.2005 | 0.0280 |
| PV1 (L2/3/4) | 0.2390 | 0.0250 | 0.2650 | 0.2505 | 0.0340 | 0.4535 | 0.0075 |
| PV2 (L5/6) | 0.0405 | 0.0670 | 0.0505 | 0.0410 | 0.1202 | 0.0075 | 0.2368 |
| SOM1(L2/3/4) | 0.1325 | 0.0250 | 0.3355 | 0.0400 | 0.0075 | 0.0660 | 0.0057 |
| SOM2 (L5/6) | 0.0510 | 0.0508 | 0.0640 | 0.0075 | 0.0425 | 0 | 0.0325 |

**Table S1B.** Synaptic strengths (unitary PSP [mV]) after averaging across layers.

| To \ From | E1 | E2 | E3 | PV1 | PV2 | SOM1 | SOM2 |
| --- | --- | --- | --- | --- | --- | --- | --- |
| E1 (L2/3) | 0.3600 | 0.2350 | 0.7800 | 0.5200 | 0.4050 | 0.3050 | 0.1250 |
| E2 (L5/6) | 0.3700 | 0.5800 | 0.7950 | 0.2325 | 0.8100 | 0.1250 | 0.2700 |
| E3 (L4) | 0.3400 | 0.1900 | 0.8300 | 0.6000 | 0.4050 | 0.2950 | 0.1400 |
| PV1 (L2/3/4) | 1.4400 | 0.6250 | 1.3400 | 0.6800 | 0.3625 | 0.5000 | 0.1125 |
| PV2 (L5/6) | 0.6600 | 2.5000 | 0.6250 | 0.4325 | 1.1900 | 0.1125 | 0.4000 |
| SOM1(L2/3/4) | 0.7750 | 0.2600 | 0.6000 | 0.4200 | 0.1050 | 0.1500 | 0.1675 |
| SOM2 (L5/6) | 0.2650 | 0.5200 | 0.2600 | 0.1050 | 0.4100 | 0.0700 | 0.4000 |

**Table S1C.**
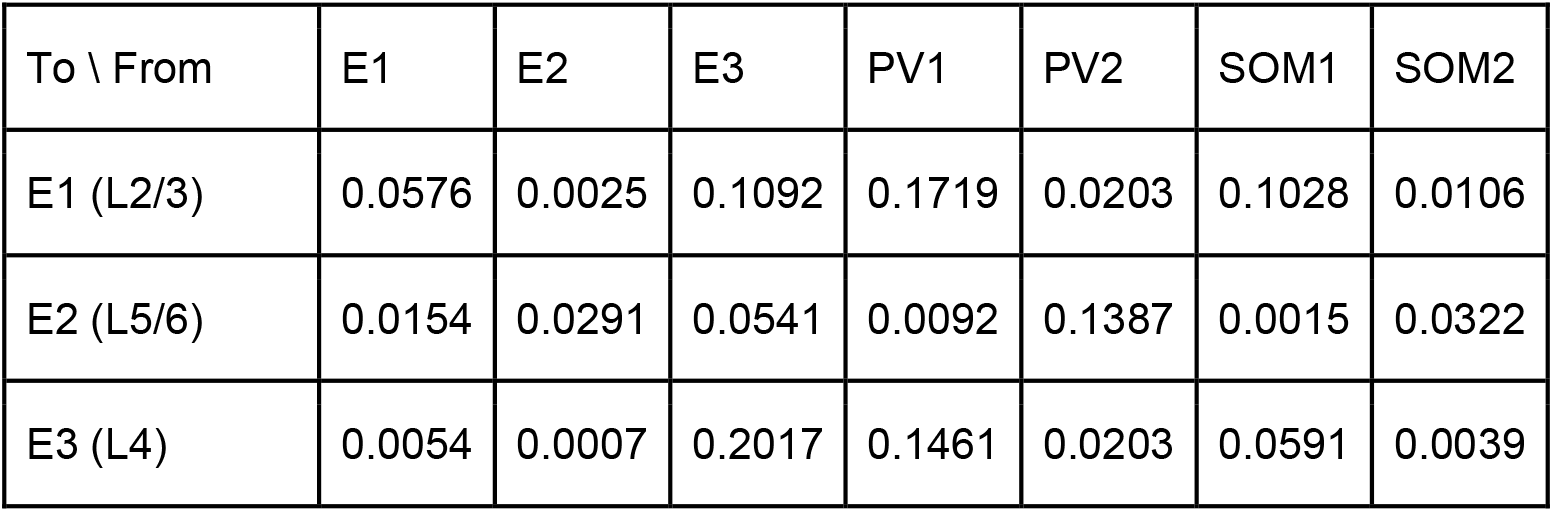
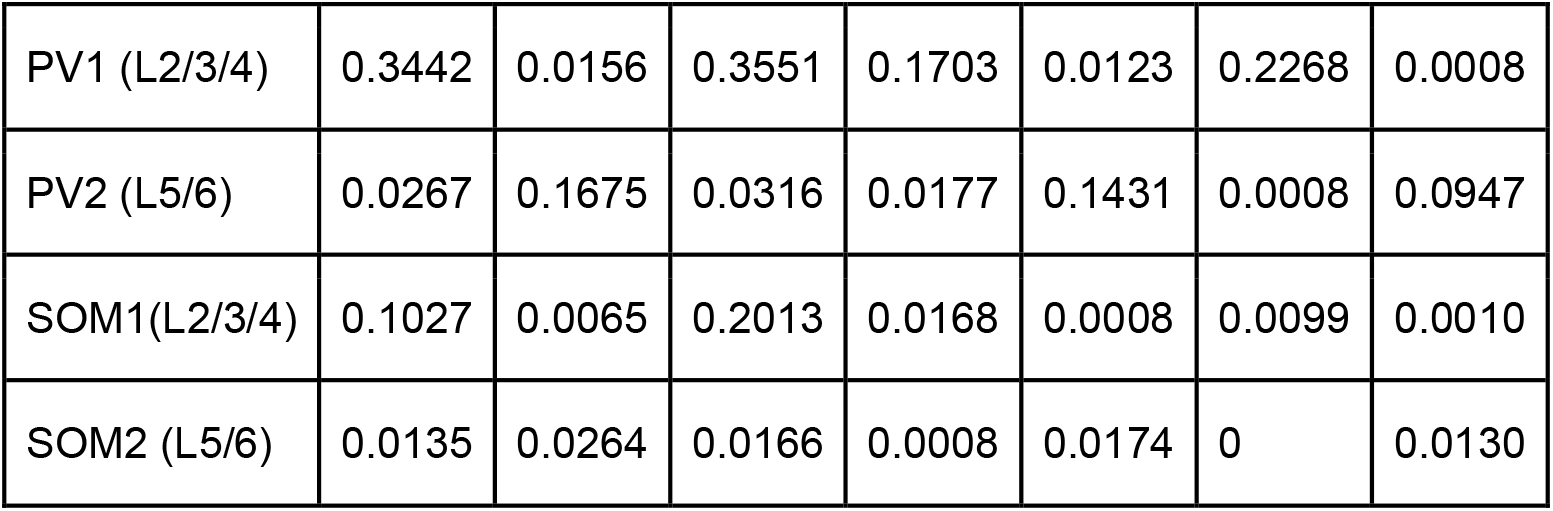
Intra-column connectivity (connection probability*synaptic strength).

